# Sca-1 expression depicts pro-inflammatory murine neutrophils under steady state and pathological conditions

**DOI:** 10.1101/2024.09.16.613221

**Authors:** Milind Nahiyera, Supriya Sinha, Priyanka Dhankani, Apurwa Singhal, Abhinav Singh, Rakesh Kumar Sharma, Ambalika Gond, Kanchan Gupta, Kalyan mitra, Amit Lahiri, Kumarvelu Jagavelu, Marie-Dominique Filippi, Madhu Dikshit, Sachin Kumar

**Author notes:** **Correspondence to:** Sachin Kumar, Pharmacology Division, Central Drug Research Institute-CSIR, Sector 10, Jankipuram extension, Lucknow-226031 (India).

## Abstract

Neutrophils play a crucial role in various pathophysiological conditions, yet targeting them for therapeutic intervention has been discouraged due to the associated risk of infections. Thus, identification of neutrophil subsets and their involvement in inflammatory conditions is warranted for targeted therapeutic strategies. This study, through screening of surface proteins on neutrophils isolated from different tissue microenvironments, identified a distinct neutrophil subset, CD11b^+^Ly6G^+^Sca1^+^ neutrophils, expressing Stem cell antigen-1 (Sca-1). Interestingly, these Sca1^pos^ neutrophils were more abundant in the liver than BM, blood, and lungs. Further analysis revealed that Sca1pos neutrophils are mature and activated with enhanced effector functions, including superoxide generation, phagocytosis, degranulation, and NETosis. Tracing studies demonstrated ageing-independent characteristics of Sca1^pos^ neutrophils. Remarkably, Sca1^pos^ pro-inflammatory neutrophils promote T cell proliferation through ROS, while inhibition of Sca-1 restores T cell proliferation and ROS generation. Intriguingly, inflammatory as well as metabolic cues induce the transition of conventional neutrophils (Sca1^neg^) to Sca1^pos^ neutrophils and differentiation of progenitors (granulocyte monocyte progenitors, GMPs) into Sca1^pos^ neutrophils. Furthermore, *in vivo* models of acute inflammation, peritonitis, and chronic inflammatory condition, non-alcoholic steatohepatitis (NASH), exhibit an increase of Sca1^pos^ neutrophils at inflammatory sites, while the pharmacological approach using NAC specifically mitigates the expansion of these pro-inflammatory neutrophils. Collectively, our findings unveil a novel subset of Sca1^pos^ neutrophils with implications for inflammation.

**Significance Statement:** Neutrophilic inflammation remains the leading driver in infectious and inflammatory diseases. Targeting neutrophil populations remained un-recommended due to hampering the immunological functions of neutrophils. The heterogeneity of neutrophils provides the perspective to target altered neutrophil subsets, but subtle changes defining neutrophil subsets make it complex and ambiguous. Our study identified abundant expression of Sca1on distinct neutrophils under steady state and inflammation. Thus, we reported previously undefined Sca1^pos^ pro-inflammatory neutrophil subsets and elucidated their regulation. This study further established their involvement in acute and chronic inflammatory conditions. This understanding may further pave the way toward targeting specific neutrophil subsets in pathologies characterized by neutrophilic inflammation.

**Graphical Abstract:** 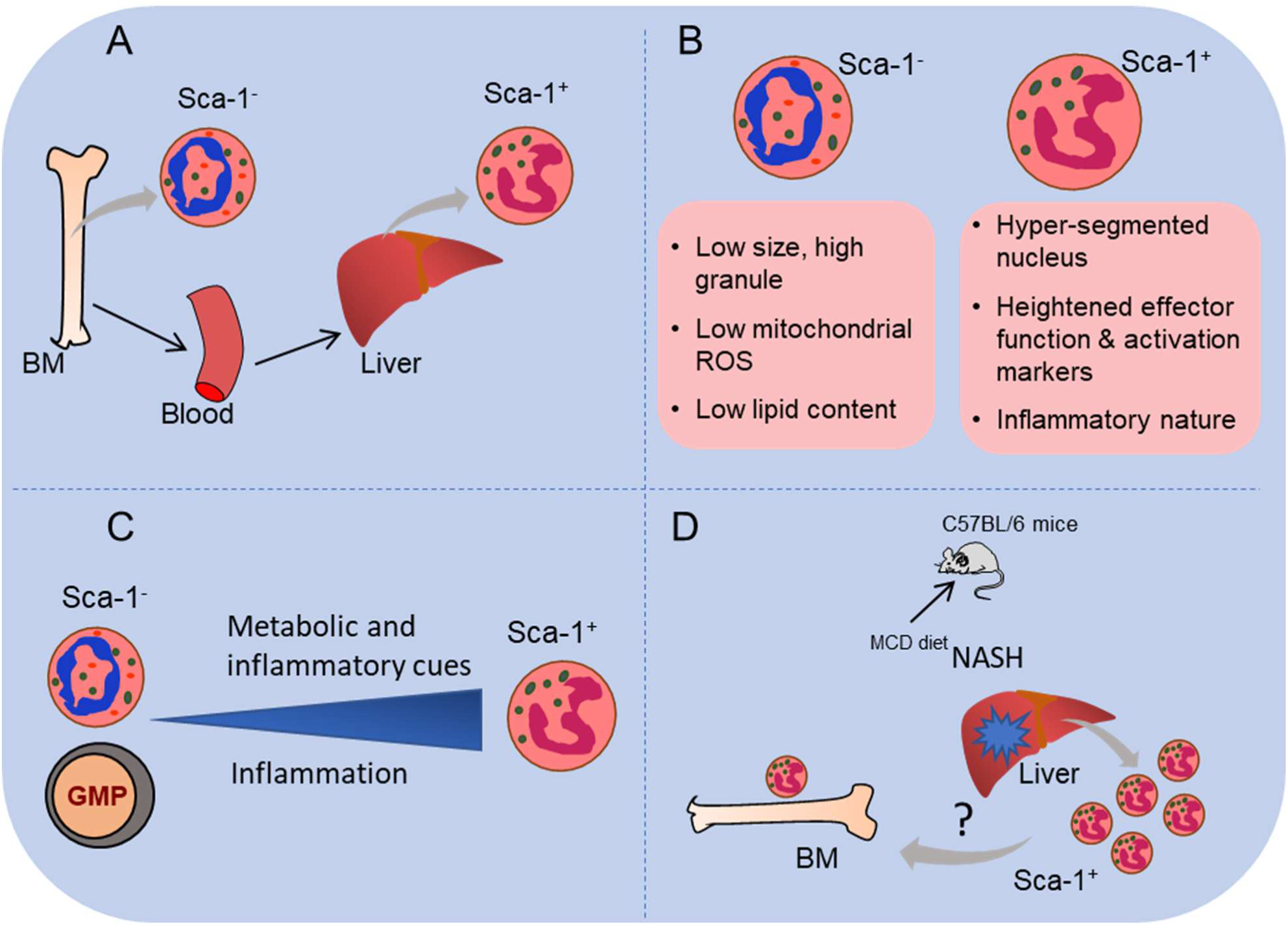

**Highlights:**

- CD11b^+^Ly6G^+^Sca1^+^ neutrophil subset identified with <1% presence in BM and >40% frequency in the liver.
- These neutrophils are mature and activated, demonstrating enhanced effector functions.
- Sca1^pos^ neutrophils promote T cell proliferation and display pro-inflammatory characteristics.
- Conventional Sca1^neg^ neutrophils transition into Sca1^pos^ neutrophils in response to inflammatory signals, while progenitors undergo differentiation.
- Both acute and chronic inflammatory models show the expansion of CD11b^+^Ly6G^+^Sca1^+^ neutrophils.

## Introduction

Neutrophils, the most abundant leukocytes, are pivotal immune system components in pathophysiological contexts. They combat invading pathogens through mechanisms including phagocytosis, oxidative burst, and the formation of neutrophil extracellular traps (NETs) (1, 2). Furthermore, neutrophils play a crucial role in intercellular communication within the immune system by secreting cytokines and chemokines to coordinate with other immune cells (3). Despite their beneficial roles, neutrophils, armed with proteases and oxidants, can also inadvertently cause tissue damage and are implicated in various pathological conditions such as sepsis, systemic lupus erythematosus (SLE), rheumatoid arthritis (RA), and systemic vasculitis (1–3). The neutrophil-lymphocyte ratio (NLR) serves as a common indicator of disease severity (1, 4). However, caution is warranted in targeting neutrophil populations despite their involvement in disease pathology, as their deficiency can lead to immunodeficiency and recurrent infections (2, 5). Recent research has shed light on neutrophil populations with diverse functions, ranging from pro-inflammatory to immunosuppressive, under pathological conditions (2, 6). Nevertheless, our understanding of neutrophil subtypes remains limited, especially in inflammation and disease. Therefore, identifying novel neutrophil subsets and elucidating their roles in inflammatory conditions is imperative. Targeting specific neutrophil subsets holds promise in providing tailored treatments for pathologies characterized by distinct neutrophil phenotypes, such as those with inflammatory or immunosuppressive features.

Neutrophils, originating in the bone marrow (BM), have a short lifespan in blood, ranging from 8 h to 5 days (7). Upon migrating into tissues, they undergo ageing and are cleared through efferocytosis (3). Throughout their lifespan, neutrophils encounter diverse tissue environments, leading to changes in density, morphology, nuclear segmentation, granular content, and gene expression. The hyperactive nature and rapid turnover of neutrophils on a circadian basis have long hindered the recognition of neutrophil subsets (8, 9). In addition, neutrophils exhibit changes in their properties and can reprogrammed in tissue microenvironment (10, 11). Advancements in technology have facilitated the identification of distinct neutrophil subtypes based on surface markers and functional responses. These subsets include aged neutrophils (CXCR4^hi^CD62L^lo^CD11b^hi^), suppressor neutrophils or granulocytic myeloid-derived suppressor cells (G-MDSCs; CD11b^+^Ly6C^low^Ly6G^+^), pro-angiogenic neutrophils (CXCR4^+^VEGFR1^+^CD49d^+^), and possibly regulatory neutrophil subtypes (12–21). In human circulation, a mature neutrophil subset characterized by CD62L^dim^CD11b^bright^CD16^bright^ has been shown to suppress T cell proliferation, increasing its activity during systemic inflammation (18). Conversely, CD16^low^CD62L^low^ cells represent an immature neutrophil population released from BM (18). CD10^+^ mature and CD10^-^ immature neutrophil subsets identified in the circulation of granulocyte colony-stimulating factor (G-CSF) treated patients with immunosuppressive characteristics. Where as in healthy individuals, CD10^+^ neutrophils promote T cell proliferation (22). Other subsets, such as OLFM-4^+^ and CD177^+^ neutrophils found in inflamed human tissues, exhibit distinct NETs and functional abilities (23, 24). In mice, neutrophils within tumor environments, known as tumor-associated neutrophils (TAN), display both anti-tumor (N1) and pro-tumor (N2) activities in a TGF-β-dependent manner (16). However, the expression levels of commonly used surface markers such as CD62L, CD11b, and CXCR4 show minimal changes, making the classification of novel neutrophil subsets complex and ambiguous (25, 26).

The present study explored the hypothesis that neutrophils exhibit phenotypic and functional diversity under distinct microenvironments. We examined neutrophils from - BM, blood, spleen, lung, and liver tissues and discovered a subset of murine neutrophils expressing Stem cell antigen-1 (Sca-1 or Ly6A/E), marked by CD11b^+^Ly6G^+^Sca1^+^, which were rare in BM (<1%) but highly abundant in liver tissue (>40%). These Sca-1 expressing neutrophils exhibited an inflammatory phenotype, displaying enhanced degranulation, phagocytosis, and other functions. Sca-1^neg^ neutrophils and granulocyte-macrophage progenitors (GMPs) could transition into the Sca-1^pos^ subset in response to inflammatory cues and metabolic signals. Additionally, the frequency of these Sca-1^pos^ neutrophils increased in inflammatory conditions such as peritonitis and liver steatohepatitis. Our study unveils and characterizes an inflammatory subset, Sca-1^pos^ neutrophils, under both steady-state and disease conditions.

## Results

### Neutrophils present in tissue microenvironments exhibit distinct characteristics

After maturation in the bone marrow, neutrophils migrate into circulation and subsequently to various tissues, a process influenced by both homeostatic and inflammatory signals (10) and potentially affecting the phenotypes and functions of neutrophils (10, 27). To explore this, we compared the phenotypic and functional profiles of neutrophils from different tissues with those from the BM. Utilizing flow cytometry, we analyzed single-cell suspensions and enriched fractions of leukocytes from the BM, blood, spleen, lung, and liver, focusing on the myeloid marker CD11b and the neutrophil marker Ly6G (**Fig 1A and Fig S1A**). Distinct percentages of neutrophils (CD11b^+^Ly6G^+^) were observed in different tissues. As foreseen, the highest abundance of neutrophils was found in the BM (60%) and blood (20%), while lower percentages (1-10%) were present in distal tissues such as the lung, spleen, and liver (**Fig 1B**). However, despite this variation in frequency, neutrophils in all tissues exhibited positivity for CD11b and Ly6G markers, with increased expression observed in the lung and liver tissues (**Fig 1A and Fig S1B, C**). This suggests potential phenotypic alterations in neutrophils within tissue microenvironments. Further analysis using flow cytometry revealed higher granularity of neutrophils in the lung and liver tissues, as indicated by increased side scatter area (SSC) (**Fig S1D**). However, no major changes in neutrophil size were observed (**Fig S1E**). The upregulation of CD11b and Ly6G along with increased SSC in distal tissues, despite lower neutrophil percentages, prompted investigations into the morphological differences of tissue neutrophils. Giemsa staining analysis of nuclear morphology revealed a ring-shaped nucleus in approximately 80% of neutrophils in the BM and blood, while 50-70% of neutrophils in the lung and liver exhibited hyper-segmented nuclei (**Fig 1C, and Fig S1F**). Together, these findings suggest alterations in surface markers, nuclear morphology, and granularity of tissue neutrophils, which may correlate with distinct characteristics and functions.

**Figure 1:**
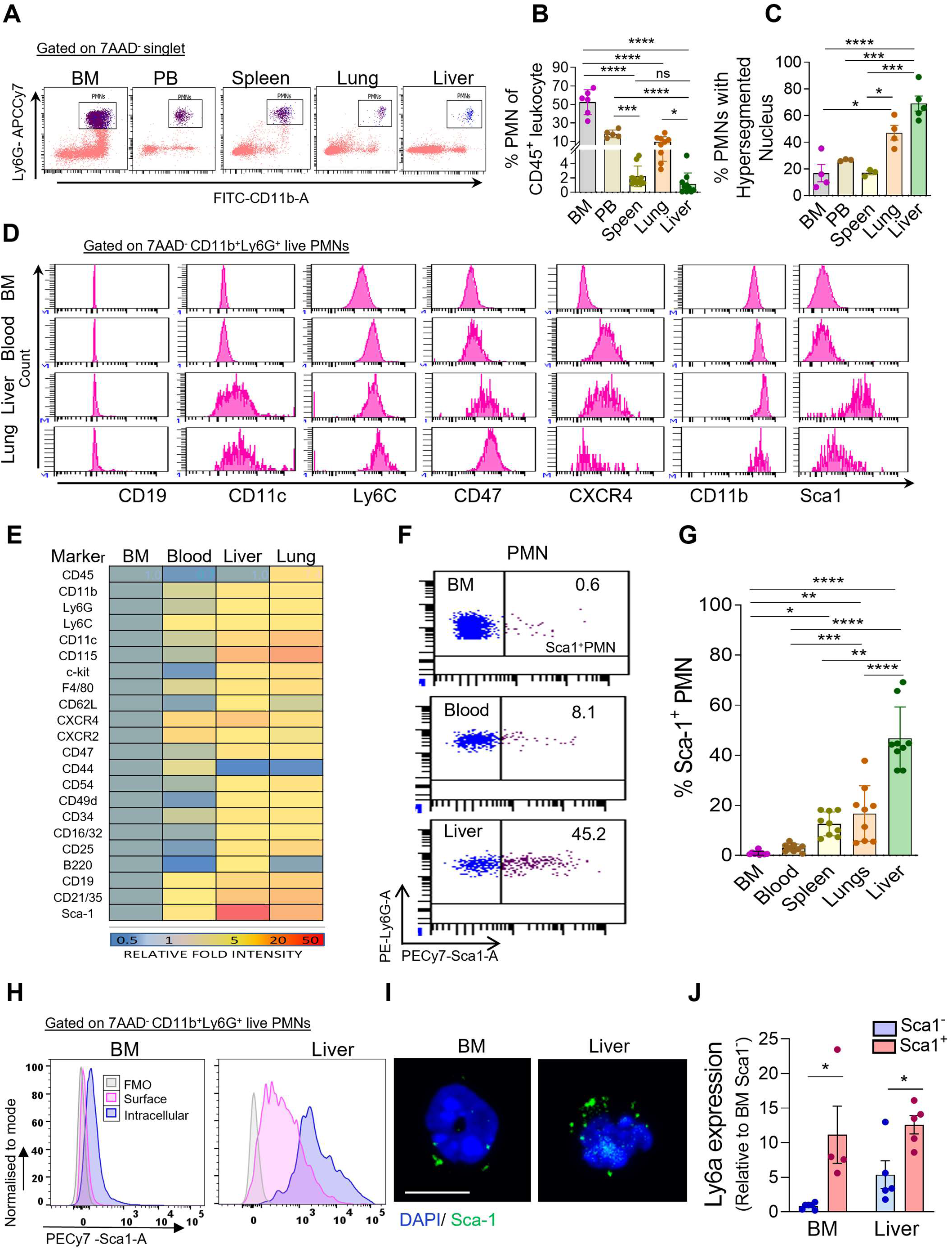
Identification of novel CD11b^+^Ly6G^+^Sca1^+^ neutrophil subset and its frequency in different tissues. **A.** Dot plot representative of flow cytometric analysis of CD11b^+^Ly6G^+^ neutrophils in different tissues. Data represent n=5 biological replicates. **B.** Bar graphs showing the percentage of neutrophils in different tissues (n=6-8 biological replicates; ns=non-significant, *p<0.05, ***p<0.001 and ****p<0.0001 using One-way ANOVA test). **C.** Quantification of the percentage of neutrophils with hyper-segmented nucleus (n=3-4 biological replicates; ns= non-significant, *p<0.05, ***p<0.001 and ****p<0.0001 using One-way ANOVA test). **D.** Histograms display the expression of various surface markers on different tissue neutrophils. Data represents n=4 biological replicates. **E.** Tabular data represents the relative expression of surface markers on tissue neutrophils with respect to BM neutrophils. Relative fold intensity is marked with a colour gradient at the bottom. Data are representative of n=4 biological replicates. **F.** Fow cytometric representative of %Sca1^+^ neutrophils in different tissue, Data represent n=8 biological replicates. **G.** Quantification of flow cytometry analysis of percentages of neutrophils exhibiting Sca1 expression in different tissues. n=9 biological replicates. *p<0.05, **p<0.01, ***p<0.001, ****p<0.0001 by One-way ANOVA test) **H.** Histogram overlay of intracellular vs. surface Sca1 expression on BM and liver neutrophils, Data are representative of n=3 biological replicates. **I.** The confocal images show the intracellular staining of Sca1 in BM and liver neutrophils. The data represent n=2 biological replicates with a minimal of 40 cells analyzed in individual experiments. Scale Bar = 5µm. **J.** Ly6a gene expression in Sca1^-^ and Sca1^+^ neutrophils sorted from BM and liver (n=4-5 biological replicates *p<0.05, by Student’s paired t-test) All data are mean ± SE.

### Identification of a distinct CD11b^+^Ly6G^+^Sca1^+^ neutrophil subset in peripheral tissues

To gain insights, we employed multicolor flow cytometry to explore the expression of various surface markers on neutrophils. Screening several candidate surface proteins revealed differential expression patterns between tissue neutrophils and those from the BM (**Fig 1D, E**). Specifically, we focused on significant changes in surface marker expression between tissue and BM neutrophils, which highlighted a striking difference in Sca-1 (Ly6A/E) expression (**Fig 1D, E**). Notably, Sca-1 was nearly absent on BM neutrophils (with <1% Sca1^pos^ neutrophils), while it was consistently more abundant on liver neutrophils (>40% Sca1^pos^ neutrophils) (**Fig 1F**). Neutrophils from blood, spleen, and lung exhibited intermediate levels, with 10-30% Sca1^pos^ neutrophils (**Fig 1F, G**). This discovery led to the identification of a previously undefined and distinct neutrophil subset expressing Stem cell antigen, denoted as CD11b^+^Ly6G^+^Sca1^+^ under steady state. It is essential to mention here that a previous study has shown Sca1 expression emerging on CD11b^+^Gr1^+^ myeloid cells of a monocytic nature during bacterial infections, but not on neutrophils (28). Furthermore, our study highlights the presence of Sca1^pos^ neutrophil subset under homeostatic states. Thus, we validated the existence of Sca1^pos^ neutrophils by removing any minimal contamination from other lineages, including lymphocytes (CD11b^neg^), eosinophils (SiglecF^pos^), and monocytic cells (Ly6C^hi^CD115^pos^). Indeed, the frequency of Sca1 expression on highly purified neutrophils (identified as LiveCD45^+^CD11b^+^Gr1^hi^Ly6G^hi^Ly6C^lo^ SiglecF^neg^CD115^-^ cells) was <1% in BM and >40% in the liver, consistent with the data observed with CD11b^+^Ly6G^+^ neutrophils (**Fig S1G**). We further established Sca1 expression on two allelic variant specific mouse stains, C57BL/6 and BALB/c, presenting *Ly6a* and *Ly6e,* respectively, and confirmed Sca1 (Ly6A/E antibody) positive neutrophils in analyzed tissues in both strains (**Fig S2A-C**). Moreover, in liver tissue, we observed Sca-1 expression on monocytic cells as well as on lymphocyte that was significantly lower than neutrophils (**Fig S2D**). Notably, experiments in which neutrophils from the different tissues were exposed to the same isolation method resulted in similar CD11b and Ly6G expression on BM and blood neutrophils, indicating tissue processing and isolation did not cause any artifacts (**Fig S2E-F**), similar observations have been recorded by others (11). Moreover, BM and blood neutrophils did not display any changes in Sca1 expression upon the same isolation processing as liver neutrophils (**Fig S2G**). These data suggest that Sca-1 expression on liver neutrophils is independent of the isolation procedure, including the collagenase treatment. We further looked into the context of tissue-resident neutrophils in the liver, where leukocytes are often found in intravascular space retained on the endothelial surface (30). Analysis of neutrophils labelled with intravenously-delivered antibody (30), confirmed more neutrophils in intravascular space in the liver **(Fig S2H)**. Further, more Sca1 expression was observed on tissue-resident neutrophils compared to intravascular neutrophils in the liver **(Fig S2I)**, suggesting tissue microenvironment-dependent regulation of Sca1 expression on neutrophils.

These findings demonstrate the existence of the Sca1^pos^ neutrophil subset, which is significantly enriched in liver tissue. Consequently, we focused our subsequent experiments on comparing Sca1^pos^ neutrophils with their Sca1^neg^ counterparts in both BM and liver tissue. To delve deeper into Sca1 expression on neutrophils, we analyzed both surface and intracellular Sca-1 levels. Intriguingly, BM neutrophils, typically displaying a Sca1^neg^ surface profile, exhibited intracellular staining of Sca1 to some extent (**Fig 1H, S2J**). Similarly, liver neutrophils displayed more abundant intracellular Sca1 expression than surface staining (**Fig 1H, S2J**). Confocal microscopy confirmed intracellular Sca1 expression in BM and liver neutrophils (**Fig 1I**). Additionally, transcript analysis of Ly6a in neutrophils corroborated Sca-1 expression in Sca1^pos^ neutrophils from both BM and liver tissue (**Fig 1J**). These comprehensive analyses across multicolor flow cytometry, intracellular staining, confocal microscopy, and transcript analysis unequivocally affirmed the identity of Sca1^pos^ neutrophils as a bona fide neutrophil subset.

### Sca1^pos^ neutrophils contain a hyper-segmented nucleus and exhibit an activated phenotype

Liver neutrophils, characterized by a notably higher percentage of Sca1 expressing cells, frequently exhibited a hyper-segmented nucleus and increased SSC (**Fig 1C, S1D**), suggesting the likelihood of Sca1^pos^ neutrophils being hyper-segmented. Indeed, analyses of sorted Sca1^pos^ neutrophils revealed larger size, heightened granularity, and hyper-segmented nuclei within the Sca1^pos^ subset compared to its Sca1^neg^ counterpart (**Fig 2A-C and S3A, B**). This pattern remained consistent across various tissues, including BM, blood, and liver. Remarkably, in BM, over 85% of sorted CD11b^+^Ly6G^+^Sca1^+^ neutrophils exhibited a hyper-segmented nucleus (**Fig 2A, B**), while BM neutrophils (predominantly Sca1^neg^) and sorted Sca1^neg^ neutrophils displayed a ring-shaped nucleus (**Fig 1C, S1F and Fig 2A, B**). Sca1^neg^ liver neutrophils also demonstrated less pronounced hyper-segmentation (**Fig 2A, B**). To further delineate the properties of Sca1^pos^ neutrophils, we scrutinized the expression of various surface markers, including CD11b, CD62L, CD54, and CXCR4, among others, on Sca1^neg/pos^ neutrophil subsets. Intriguingly, liver neutrophils exhibited an up-regulation of CD11b, Ly6G, CXCR4, CD47, CD54, and CD49d surface expression (**Fig S1B, C and S3C**), a pattern further confirmed by Sca1^pos^ neutrophils across all analyzed tissues (**Fig 2D, E and Fig S3D**). Moreover, heightened expression of surface proteins associated with cell activation, adhesion, and migration (CD11b, CD49d, CD54), along with the “don’t eat me” signal (CD47), suggested an activated state of the Sca1^pos^ neutrophil subset (**Fig 2D, E and Fig S3C**). Notably, levels of selectin CD62L were comparable between Sca1^pos^ and Sca1^neg^ neutrophil subsets across BM, blood, and liver tissue (**Fig 2F, Fig S3C**). These findings underscore structural and phenotypic alterations in Sca1^pos^ neutrophils.

**Figure 2:**
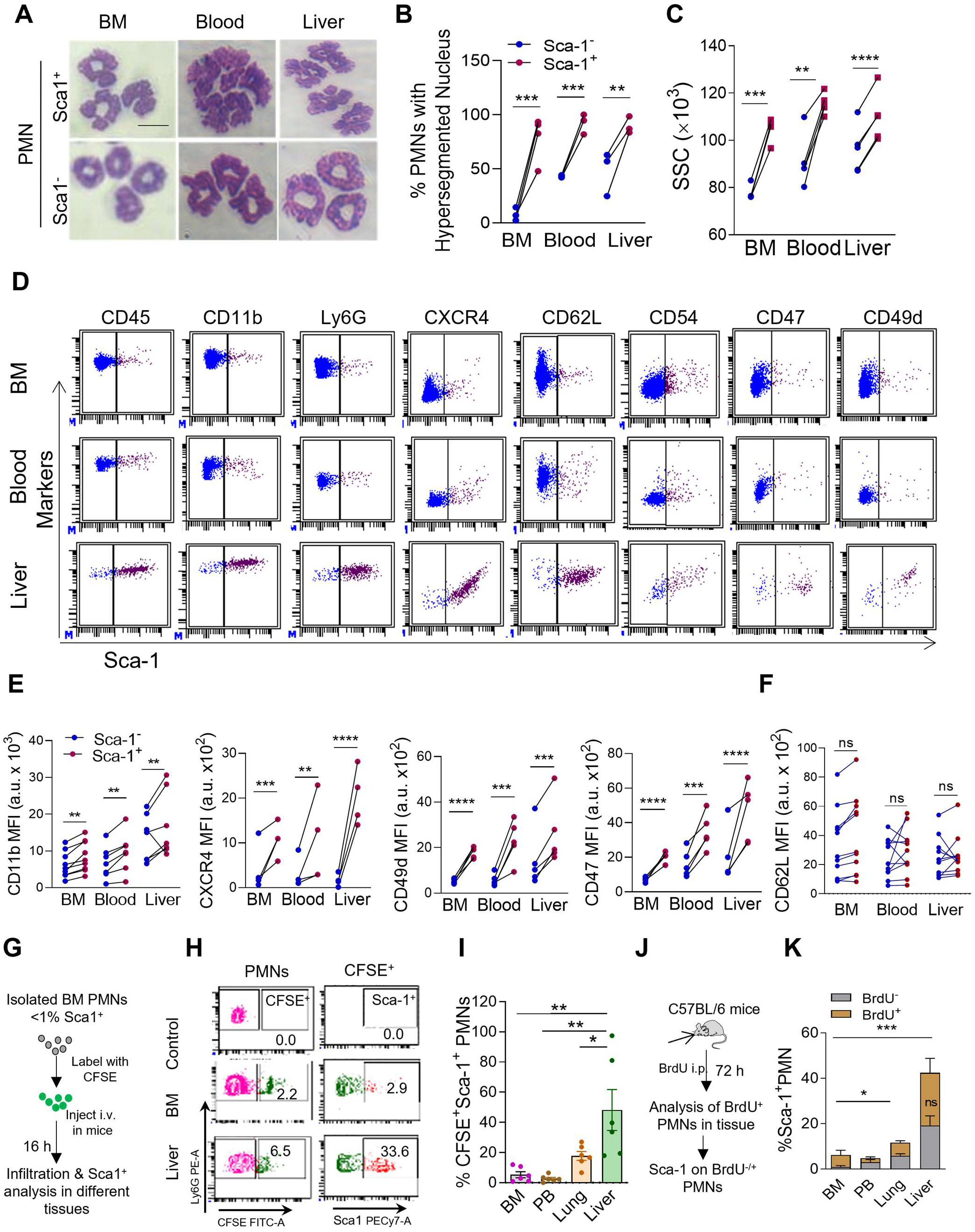
Sca1^+^ neutrophils are active neutrophils but do not present aged neutrophils. **A.** Representative of Giemsa staining of Sca1^-^ and Sca1^+^ neutrophils sorted from BM, blood, and liver. Data represent n=3 biological replicates. Scale Bar = 5µm. **B.** Quantification of hyper-segmented nuclei in Sca1^-^ and Sca1^+^ neutrophils sorted from BM, blood and liver (n=3 biological replicates **p<0.01, ***p<0.001 using Student’s paired t-test) **C.** Quantification of SSC of Sca1^-^ and Sca1^+^ neutrophils of BM, blood, and Liver performed using flow cytometry (n=3 biological replicates. **p<0.01, ***p<0.001 using Student’s paired t-test) **D.** Flow cytometry dot plot representing the surface expression of proteins related to cellular activation, adhesion, and migration, including CD45, CD11b, Ly6G, CXCR4, CD62L, CD54, CD47, CD49d on Sca1^-^ and Sca1^+^ neutrophils of BM, blood, and liver. Data represent n=3 minimal biological replicates. **E.** Quantification of CD11b, CXCR4, CD49d, and CD47 surface expression on Sca1^-^ and Sca1^+^ neutrophils of BM, blood, and liver. (n=3-5 biological replicates. **p<0.01, ***p<0.001 using Student’s paired t-test). **F.** Flow cytometry quantification of CD62L surface expression of Sca1^-^ and Sca1^+^ neutrophils n=5 biological replicates. ns= non-significant by Student’s paired t-test **G.** Experimental plan for tracing of CFSE labeled adoptive transferred BM neutrophils (<1% Sca1^+^) via i.v. injection. **H.** Flow cytometric representative of plot shows % CFSE^+^ neutrophils and % CFSE^+^ Sca1^+^ neutrophils in BM and Liver. Data represent n=6 biological replicates. **I.** Quantification of %CFSE^+^Sca1^+^ neutrophils in BM and Liver n=6 biological replicates. *p<0.05, **p<0.01 by One-way ANOVA test). **J.** Experiment plan of in vivo BrdU labeling of Sca1^-^ and Sca1^+^ neutrophils **K.** Quantification of Sca1^+^ neutrophils in BrdU^-/+^ neutrophils *p<0.05, ***p<0.001 between Sca1^+^ neutrophils in different tissue and ns= non-significant between Sca1^-^ and Sca1^+^ neutrophils in liver tissue by two-way ANOVA) All data are mean ± SE.

### Sca1^pos^ neutrophils display ageing-independent regulation and characteristics

We next evaluated whether Sca1 expression on neutrophils is linked to ageing? Interestingly, other markers commonly associated with ageing, such as CD11b and CXCR4, were notably heightened on Sca1^pos^ neutrophils. To understand potential relationship between ageing and Sca1 expression on neutrophils, CFSE-labeled BM neutrophils (with <1% Sca1^pos^ neutrophils) were adoptively transferred and tracked in various tissues, including BM, blood, lung, and liver (**Fig 2G**). Analysis of live CD11b^+^Ly6G^+^ neutrophils unveiled 1-10% of labeled CFSE^+^ neutrophils in different tissues (**Fig 2H, S3E**). Surprisingly, 30% of these labeled neutrophils in the liver exhibited a Sca1^pos^ status, contrasting with ∼3% in BM (**Fig 2H, I**). Host tissue (unlabeled) neutrophils displayed the anticipated Sca1 expression pattern of <1% in BM and ∼40% in liver (**Fig S3F**). This distinct level of Sca1 expression on adoptively transferred similar-age BM neutrophils implies that Sca1 expression on neutrophils is independent of ageing but likely influenced by the tissue microenvironment and other factors.

Furthermore, BrdU labeling of neutrophils also confirmed the lack of correlation between Sca1 expression and BrdU staining, as BrdU^neg/pos^ neutrophils exhibited comparable Sca1 expression **(Fig 2J, K and S3G)**. Consistent to data so far, liver tissue neutrophils displayed higher Sca1 expression compared to other analyzed tissues, independent of BrdU staining **(Fig 2J, K and S3G)**. Of note, neutrophils in blood, lung, and liver exhibited approximately 50% BrdU positivity, whereas BM contained fewer than 10% BrdU^+^ neutrophils, likely due to the egress of newly formed BrdU^+^ neutrophils post 72h **(Fig S3G, H)**. These findings confirm that Sca1 expression on neutrophils is not directly associated with the age of these cells.

### Sca1^pos^ neutrophil subset exhibits enhanced effector functions

Neutrophils display versatile functions, including phagocytosis, reactive oxygen species (ROS) generation, and others for pathogen elimination (31). We analysed these effector functions in Sca1^pos^ and Sca1^neg^ neutrophil subsets. Intriguingly, Sca1^pos^ neutrophil subset exhibited elevated levels of cellular and mitochondrial superoxide (MitoSOX) compared to Sca1^neg^ neutrophils (**Fig 3A, B**). Furthermore, mitochondrial membrane potential, as indicated by MitoTracker Red, was enhanced in the Sca1^pos^ neutrophil subset (**Fig 3C**). More mitochondrial activity has been previously observed in the inflammatory neutrophil subsets (21, 32). Additionally, Sca1^pos^ neutrophils presented significantly higher levels of nitric oxide (NO) and CD54 (ICAM) compared to Sca1^neg^ neutrophils (**Fig 3D, E**). These data are irrespective of their location (Fig 4B/C) and location, imparting their phenotype, particularly in the liver (**Fig S4A-D**). This might depend on the association of function with Sca1^pos^ neutrophils or the milieu exposed to regulating specific function, which needs further investigation, including the possibility of a maturation process happening in the liver. Functional analysis identified the heightened phagocytic potential of the Sca1^pos^ neutrophil subset compared to its Sca1^neg^ counterpart (**Fig 3F, G, S4E-H**), contributing to the overall higher phagocytic activity of liver neutrophils compared to BM neutrophils (**Fig S4E, F**) (26). This underscores liver neutrophils’ enhanced functionality and active state, primarily driven by the Sca1^pos^ subset. Moreover, analysis of intracellular cytokines, including IL6 and IL-1β, also indicated higher expression in Sca1^pos^ neutrophils (**Fig 3 H, I**).

**Figure 3:**
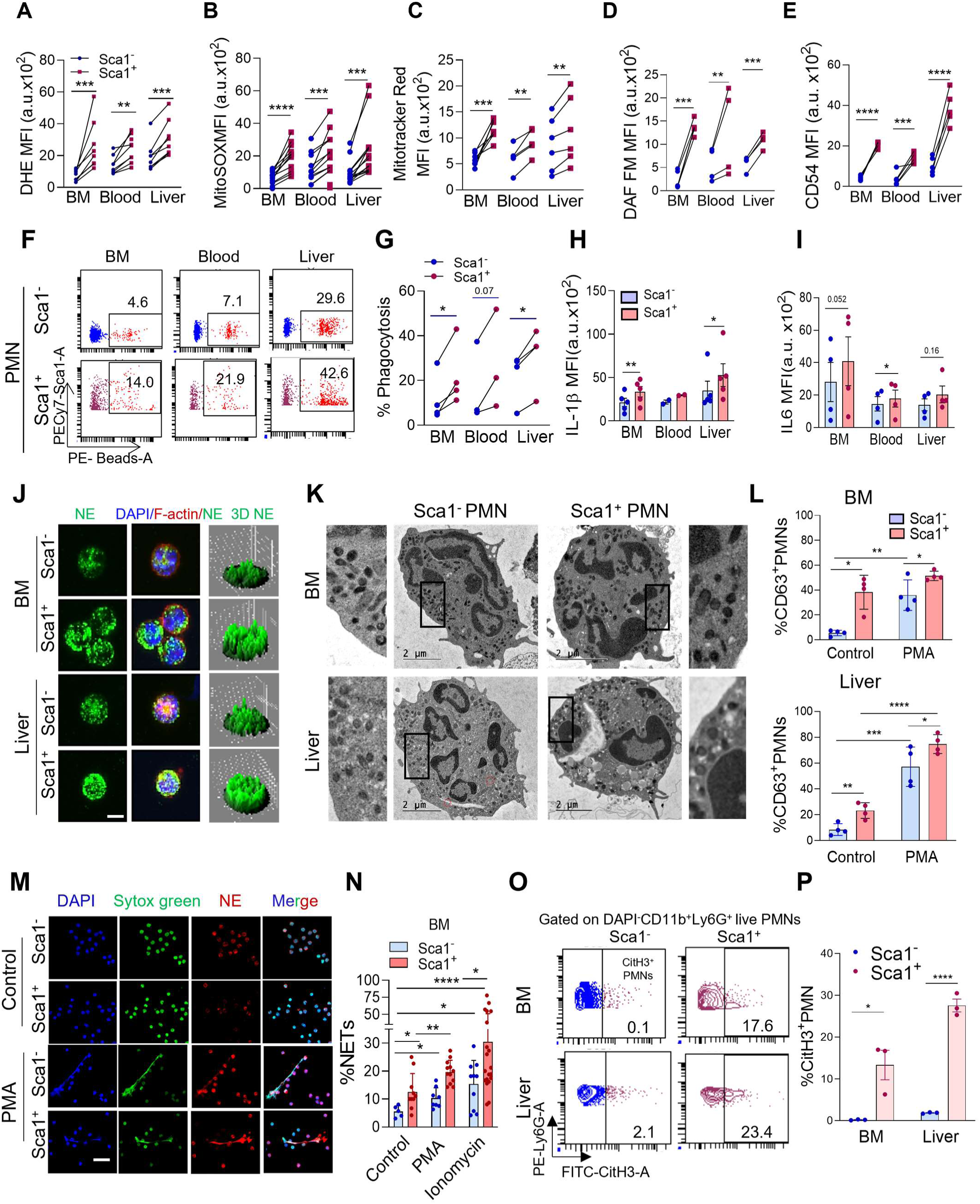
Sca1^+^ neutrophils exhibit more neutrophil effector function than conventional neutrophils. Sca-1^pos^ and Sca-1^neg^ neutrophils from mice BM, blood and liver tissues were either stained and analysed for functional assays or sorted to perform functional studies. **A.** Flow cytometry-based quantification of superoxide anion detected by DHE in labeled Sca1^-^ and Sca1^+^ neutrophils. n=8 biological replicates; **p<0.01, ***p<0.001, using Student’s paired t-test. **B.** Mitochondrial superoxide by MitoSox staining in labeled Sca1^-^ and Sca1^+^ neutrophils. n=12 biological replicates. ***p<0.001, ****p<0.0001, using Student’s paired t-test. **C.** Analysis of Mitotracker Red in labeled Sca1^-^ and Sca1^+^ neutrophils. n=6 biological replicates; **p<0.01, ***p<0.001, using Student’s paired t-test. **D.** Nitric oxide (NO) level detected by DAF in labeled Sca1^-^ and Sca1^+^ neutrophils. n=4 biological replicates. **p<0.01, ***p<0.001, using Student’s paired t-test. **E.** Surface expression of ICAM1 (CD54) in labeled Sca1^-^ and Sca1^+^ neutrophils. n=3-5 biological replicates. ***p<0.001, ****p<0.0001, using Student’s paired t-test. **F.** Flow cytometry dot plot representative of %phagocytosis of latex beads by Sca1^-/+^ neutrophils of BM, blood, and Liver. Data represent n=3-4 biological replicates. **G.** Flow cytometric quantification of %phagocytosis of latex beads by Sca1^-/+^ neutrophils. n=3-4 biological replicates. *p<0.05, **p<0.01, using Student’s paired t-test. **H.** Quantification of the intracellular release of IL-1β and IL-6 of Sca1^-/+^ neutrophils of BM, blood and Liver done by flow cytometry. n=2-5 biological replicates. *p<0.05, **p<0.01, using Student’s paired t-test. **J.** Confocal images of sorted BM and Liver Sca1^-/+^ neutrophils stained for elastase and F-actin; 3D plot of neutrophil elastase is provided on the right side. Data represent n=2 biological replicates. Scale Bar = 5µm. **K.** TEM images representing the structural organization with various granule content and distribution in sorted BM and Liver Sca1^-/+^ neutrophils. Data are from n=2 biological replicates with a minimal 20 cells per experiment analyzed. **L.** Flow cytometry quantification of CD63 expressing Sca1^-^ and Sca1^+^ neutrophils of BM and Liver. n=4 biological replicates. *p<0.05, **p<0.01, ***p<0.001, ****p<0.0001 using Student’s paired t-test. **M.** Immunofluorescence images of NETs induction as marked with nuclear stain DAPI and neutrophil elastase in BM Sca1^-/+^ neutrophils with or without PMA treatment. n=4 biological replicates. Scale Bar = 40µm. **N.** Quantification of %NETs induction upon PMA and ionomycin treatment by Sca1^-/+^ BM neutrophils. n=5-9 biological replicates. *p<0.05, **p<0.01, ****p<0.0001 using Student’s unpaired t-test. **O.** Contour plot represents % Sca1^-/+^ CitH3^+^ neutrophils in BM and Liver. Data are from n=3 biological replicates **P.** Flow cytometry quantification of % CitH3^+^ in BM and Liver Sca1^-/+^ neutrophils. n=3 biological replicates. *p<0.05, ****p<0.0001 using Student’s paired t-test. All data are mean ± SE.

**Figure 4:**
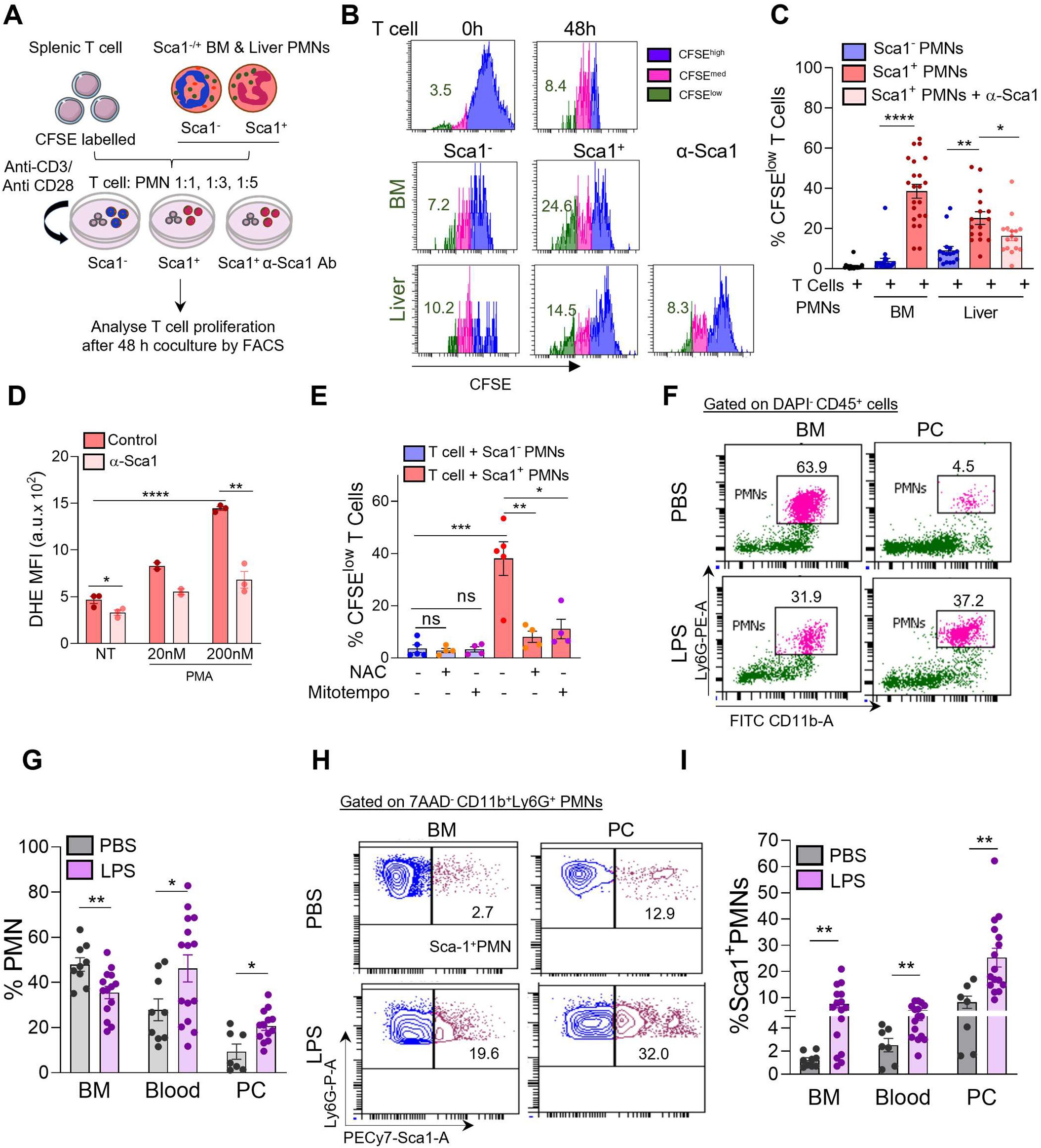
Sca1^+^ neutrophils enhanced T-cell proliferation and induced under inflammatory conditions. **A.** Experimental plan of CFSE labeled T cell proliferation with Sca1^-/+^ neutrophils **B.** Histogram shows T cell division as CFSE^low^ T cell after coculture with Sca1^-^ and Sca1^+^ neutrophils with T cell: PMN ratio (1:3). Top panel shows input 0 h and post 48 h T cell CFSE staining using histogram presentation. The right side panel presents Sca1^+^ effect on T cell proliferation in the presence of α-Sca1 Ab. **C.** Quantification of CFSE^low^ T cell presenting division after coculture with Sca1^-^ and Sca1^+^ neutrophils with T cell: PMN ratio (1:3) and in the presence of Sca1 neutralizing antibody. n=8-12 biological replicates. *p<0.05, **p<0.01, ***p<0.001, ****p<0.0001, using Student’s paired t-test. **D.** Quantification of superoxide levels in Liver neutrophils detected by DHE upon stimulation with low and high concentrations of PMA upon Sca1 blocking. n=2-3 biological replicates. *p<0.05, **p<0.01, ****p<0.0001, using Student’s paired t-test. **E.** T cell proliferation with Sca1^-^ and Sca1^+^ neutrophils treated with N Acetylcysteine (NAC) and Mito tempo. ns= nonsignificant, *p<0.05, **p<0.01, ***p<0.001 using Student’s unpaired t-test. **F.** FACS dot plot representative of %PMNs of BM and peritoneal cavity of *in vivo* peritonitis model. Data represent n=2 experiments with 6-8 mice. **G.** Quantification of %PMNs of BM, blood, and peritoneal cavity of *in vivo* peritonitis model. Data represent n=2 experiments with 6-8 mice. *p<0.05, using Student’s unpaired t-test. **H.** Flow cytometric contour plot representative of % Sca1^+^ neutrophils of BM, liver, and peritoneal cavity of *in vivo* peritonitis model. Data represent n=2 experiments with 6-8 mice. **I.** Quantification of %Sca1^+^ neutrophils of BM, liver and peritoneal cavity of *in vivo* peritonitis model. Data represent n=2 experiments with 6-8 mice. *p<0.05, **p<0.01, ***p<0.001 using Student’s unpaired t-test. All data are mean ± SE.

Phagocytosis and cytokine release often involve degranulating various neutrophil granules (2, 5). Confocal microscopy-based analysis revealed the peripheral localization of elastase-positive primary granules in Sca1^pos^ neutrophils, contrasting with Sca1^neg^ neutrophils from both BM and liver tissue (**Fig 3J**). Furthermore, Transmission Electron Microscopy (TEM) analysis demonstrated distinct pattern of granule distribution in Sca1^pos^ neutrophils, with primary granules identified based on the prominent and darker appearance (33), located mainly at the outer periphery, unlike the well-distributed granule populations observed in Sca1^neg^ BM neutrophils (**Fig 3K**). Similar observations were noted in liver Sca1^neg^ and Sca1^pos^ neutrophils (**Fig 3K**), suggesting heightened degranulation in the Sca1^pos^ neutrophil population. Elevated CD63 surface expression confirmed increased degranulation in Sca1^pos^ neutrophils, which was further augmented upon stimulation (**Fig 3L, S4I**).

Given that Sca1^pos^ neutrophils exhibit enhanced superoxide levels and granule release, which can cause damage to cellular components, including lipids, proteins, and nuclear content. We further investigated whether these cells are prone to neutrophil extracellular traps (NETs) release. Indeed, analyses revealed increased NETs released by Sca1^pos^ neutrophils under steady-state conditions and with known inducers, including PMA and ionomycin (**Fig 3M, N**). Moreover, Sca1^pos^ neutrophils displayed higher levels of CitH3, a NETs marker, with 10% and 30% positivity observed in BM and liver, respectively, while Sca1^neg^ neutrophils exhibited minimal levels (**Fig 3O, P and Fig S4J**). These observations collectively confirm the enhanced effector functions of Sca1^pos^ neutrophils, including superoxide generation, phagocytic potential, degranulation, and NET formation.

### Sca1^pos^ neutrophils exhibit an inflammatory phenotype

Exploring the functional role of this neutrophil subset, whether it leans towards a pro-inflammatory or anti-inflammatory profile, is indeed intriguing. To delve into this, CFSE-stained T cells were co-cultured with Sca1^neg/pos^ BM and liver tissue neutrophils at varying T cell: neutrophil ratios (1:1, 1:3, and 1:5), and T-cell proliferation was assessed as CFSE dilution using flow cytometry (**Fig 4A**). Surprisingly, Sca1^pos^ neutrophils induced significantly more T cell proliferation than Sca1^neg^ neutrophils (**Fig 4 B, C, S5A**). This is consistent with NETs and ROS increasing the proliferation of T cells (34–37).

Interestingly, this induced T cell proliferation was partially inhibited by α-Sca1 blocking antibody treatment (**Fig 4 B, C**). Furthermore, α-Sca1 blocking antibody mitigated superoxide generation in liver neutrophils, confirming the role of Sca1 in regulating ROS (**Fig 4D**). Treatment with N-acetylcysteine (NAC) and Mito-Tempo significantly attenuated the enhanced T cell proliferation by Sca1^pos^ neutrophils (**Fig 4E**), suggesting the involvement of neutrophil ROS in promoting T cell proliferation. To further corroborate the association of Sca1^pos^ neutrophils with inflammation, we examined Sca1^pos^ neutrophils in an LPS-induced peritonitis model (**Fig S5B**). LPS challenge resulted in the infiltration of immune cells, including neutrophils, into the peritoneal cavity (**Fig S5C**). Flow cytometry analysis revealed an increase of neutrophils in the peripheral blood and peritoneal cavity following LPS treatment and likely egress from BM in response to the inflammatory insult (**Fig 4F, G**). Importantly, Sca1^pos^ neutrophil subsets were significantly augmented upon LPS treatment in the blood, peritoneal cavity, and BM (**Fig 4H, I**), suggesting that inflammatory stimuli trigger the regulation of Sca1 expression on neutrophils. These findings provide compelling evidence confirming the pro-inflammatory nature of Sca1^pos^ neutrophils.

### Metabolic and Inflammatory cues orchestrate the expansion of Sca1^pos^ neutrophil subset

During TEM analysis of Sca1^neg/pos^ neutrophils, we made an intriguing observation of Sca1^pos^ liver neutrophils displaying accumulation of lipid droplets **(Fig 5A, B)**. This finding was further validated by Bodipy-based staining, which confirmed a higher lipid content in Sca1^pos^ neutrophils (**Fig 5C, D**). Additionally, liver neutrophils exhibited elevated lipid content compared to BM neutrophils (**Fig S5D**), and this lipid accumulation was directly correlated with Sca-1 expression on neutrophils (**Fig 5E**), suggesting regulation of Sca1 expression on neutrophils mediated by lipid metabolism. The increase in the Sca1^pos^ subset during peritonitis suggests the potential conversion of Sca1^neg^ neutrophils into Sca1^pos^ neutrophils; therefore, we delved into investigating the origin and regulation of the Sca1^pos^ neutrophil subset. Various inflammatory and metabolic cues, such as palmitic acid, LPS, and zymosan, were tested in *in-vitro* cultures with BM neutrophils (Sca1^neg^ neutrophils). Interestingly, these cues increased in the Sca1^pos^ subset, albeit remaining <10% of total neutrophils (**Fig 5F, G**).

**Figure 5:**
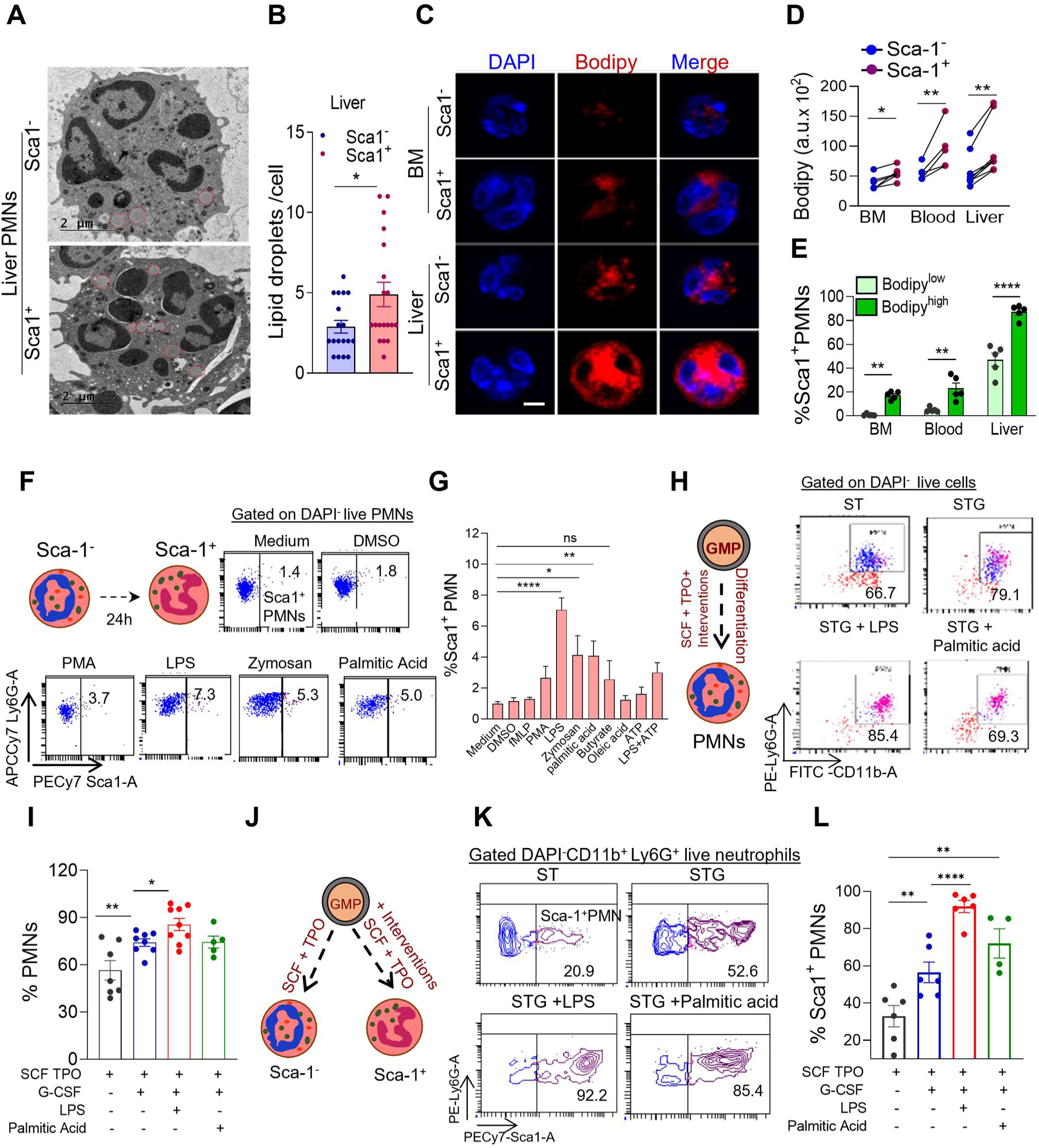
Regulation and switching of conventional neutrophils into Sca1^+^ neutrophils. **A.** TEM images of Sca1^-^ and Sca1^+^ neutrophils sorted from liver shows lipid droplets (red circle). Data are from n=2 biological replicates and >20 cells analyzed. Scale Bar = 5µm. **B.** Quantification of lipid droplets/cell of liver Sca1^-^ and Sca1^+^ neutrophils. Data represent n=2 experiments with 6-8 mice. *p<0.05, using Student’s unpaired t-test. **C.** Confocal images of Bodipy staining in sorted Sca1^-^ and Sca1^+^ neutrophils. Data represent n=3 biological replicates with at least 30 cells analyzed in Z stacks. **D.** Flow cytometric analysis of Bodipy staining in Sca1^−^ and Sca1^+^ neutrophils isolated from BM, blood and liver. Data represent n=5-8 experiments. *p<0.05, **p<0.01, ***p<0.001 using Student’s paired t-test. **E.** %Sca1^+^ neutrophils of Bodipy^low^ and Bodipy^high^ neutrophils of BM, blood, and Liver. n=5 biological replicates. *p<0.05, **p<0.01, ****p<0.0001 using Student’s paired t-test. **F.** Schematic diagram and dot plot representative *in vitro* culture of BM conventional neutrophils (>99%Sca1^-^) with various interventions and analysis towards switches to Sca1^+^ neutrophils. n=5-8 biological replicates. **G.** Flow Cytometric quantification shows %Sca1^+^ PMNs switching from *in vitro* culture BM conventional neutrophils with various interventions. n=5-8 biological replicates. *p<0.05, **p<0.01, **p<0.01, ***p<0.001 using One way ANOVA test. **H.** Schematic diagram and dot plot representing neutrophilic differentiation under *in vitro* culture of GMPs with/without inflammatory mediators. STG stands for SCF+ TPO+ G-SCF. n=7 biological replicates. **I.** Bar graph presenting Flow cytometric quantification of %PMNs differentiated from GMPs with/without mediators. n=7 biological replicates. *p<0.05, **p<0.01, **p<0.01 using Student’s paired t-test. **J.** Schematic diagram represents *in vitro* culture of GMPs with/without inflammatory mediators and putative differentiation into Sca1^-/+^ neutrophils. **K.** Contour plot represents differentiated Sca1^-/+^ neutrophils from GMPs with/without *in vitro* inflammatory mediators. Data represent n=4-6 biological replicates. **L.** FACS quantification of Sca1^+^ neutrophils differentiated from *in vitro* cultured GMPs with various inflammatory cues. n=4-6 biological replicates. *p<0.05, **p<0.01 using Student’s paired t-test. All data are mean ± SE.

This subtle increase could be attributed to the conventional differentiation of Sca1^neg^ neutrophils; hence, we explored the progenitor (GMP) to neutrophil differentiation model. GMPs cultured in the presence of stem cell factor (SCF) and Thrombopoietin (TPO) produced neutrophils within 4-6 days (38), which were further augmented with G-CSF and other treatments (**Fig 5H, I**). Remarkably, neutrophils in the SCF and TPO group exhibited a 20% Sca1^pos^ neutrophil subset. Surprisingly, this proportion was further boosted to 70-90% Sca1^pos^ neutrophils upon treatment with G-CSF, LPS, and palmitic acid (**Fig 5 J-L**). This underscores the reprogramming of progenitors and the transition of Sca1^neg^ into Sca1^pos^ neutrophils mediated by inflammatory and metabolic signals.

### Expansion of Sca1^pos^ neutrophils in non-alcoholic steatohepatitis disease model

We examined the significance of the Sca1^pos^ neutrophil subset in the chronic inflammatory condition, NASH. In contrast to the standard chow diet, mice fed a methionine-choline-deficient (MCD) diet developed steatohepatitis and inflammation (**Fig 6A**). Notably, neutrophil infiltration and activation are linked to fatty liver, NASH, and the severity of steatosis (39, 40), yet the specific neutrophil subtypes associated with these diseases remained ill-defined. The MCD diet-induced non-alcoholic steatohepatitis was confirmed by H&E and Oil Red staining (**Fig 6B**), Liver/Body weight ratio, and elevated serum levels of alanine transaminase (ALT) and aspartate aminotransferase (AST) (**Fig 6C**).

**Figure 6:**
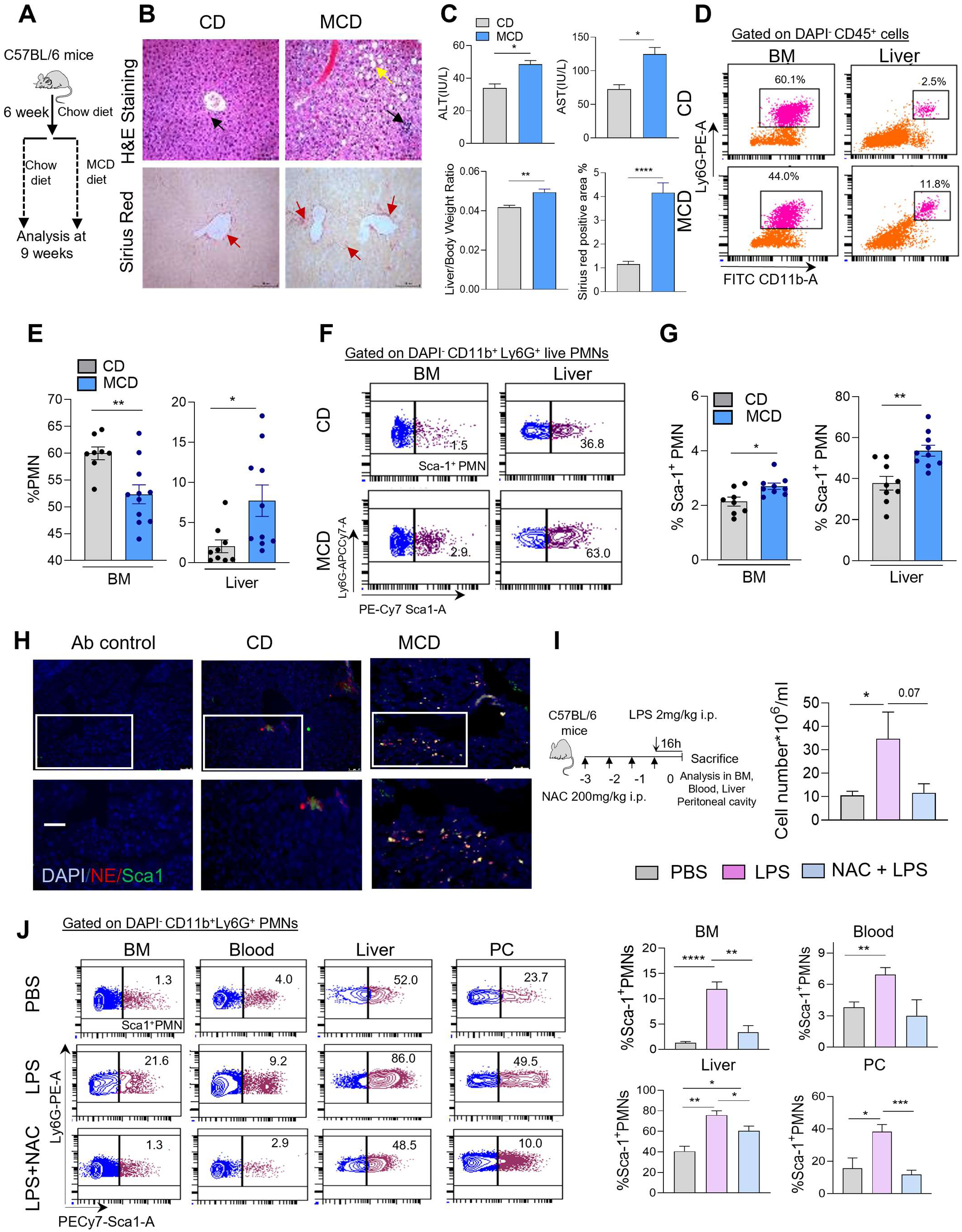
Sca1^+^ neutrophils in liver inflammatory conditions. **A.** Schematic diagram shows NASH mouse model developing strategy with Methionine and choline-deficient diet. **B.** Top panel; H&E staining shows lipid droplet formation (yellow arrow) and immune cell infiltration (black arrow) in CD and MCD fed mice liver whereas red arrow of Sirius red staining in lower panel represents collagen deposition. n=4-6 biological replicates. **C.** Quantification of biochemical analysis of serum AST, ALT, Liver/body weight ratio, and Sirius red positive area in CD and MCD-fed mice. n=4-6 biological replicates, *p<0.05, **p<0.01, ****p<0.0001 using Student’s paired t-test. **D.** Flow Cytometric dot plots represent %PMNs in BM, and liver of chow and MCD fed mice. Data represent n=2 experiments with 8-12 mice. **E.** Quantification of %PMNs in BM and liver of chow and MCD fed mice. Data represent n=2 experiments with 8-12 mice. *p<0.05, **p<0.01, using Student’s paired t-test. **F.** Flow Cytometric contour plot demonstrating %Sca1^+^ neutrophils in BM and liver of chow and MCD fed mice. Data represent n=2 experiments with 8-12 mice. **G.** Quantification for %Sca1^+^ neutrophils in BM and liver of chow and MCD fed mice. Data represent n=2 experiments with 8-12 mice. p=0.06, ***p<0.001, using Student’s paired t-test. **H.** Immunofluorescence staining of neutrophil elastase and Sca1 in liver tissue cryo-sections of CD and MCD fed mice. Data represents n=4 biological replicates. Scale Bar = 50 µm. **I.** Experiment setup of LPS-induced peritonitis model and quantification of total cell number of the peritoneal cavity in control and LPS-treated mice. n=4 biological replicates. *p<0.05 using Student’s unpaired t-test. **J.** Flow cytometry representative and quantification for % Sca1^+^ neutrophils in BM, blood, peritoneal cavity and liver of LPS induced peritonitis model with/without NAC treatment. Data represent n=2 experiments with 6-8 mice. *p<0.05, **p<0.01, ***p<0.001, ****p<0.0001 using Student’s paired t-test. All data are mean ± SE.

Flow cytometric analysis of neutrophils in chow and MCD diet-fed mice revealed increased neutrophils in the MCD-treated liver (Fig 6D, E). Further investigation showed a significant increase in Sca1^pos^ neutrophils in the liver and bone marrow tissue in MCD-fed mice (**Fig 6F, G**). Immunocytochemistry analyses of liver tissue sections also confirmed enhanced Sca1 expression in neutrophils in MCD-fed mice (**Fig 6H**). These findings suggest a potential pathological role of Sca1^pos^ neutrophils in chronic inflammatory condition, NASH. To further explore the relevance of Sca1^pos^ neutrophils in inflammation, we investigated the effect of NAC pretreatment on Sca1^pos^ neutrophils in LPS-challenged animals (**Fig 6I**). NAC rescued the infiltration of immune cells, majorly macrophages and lymphocytes (**Fig S5E**) (41). Though total PMNs were not changed in NAC treated group (**Fig S5F**) (42), NAC pre-treatment rescued pro-inflammatory Sca1^pos^ neutrophils in LPS-inflamed liver, as well as normalized their presence in bone marrow, blood, and peritoneal cavity (**Fig 6J**). These data underscore a significant association between Sca1^pos^ neutrophil subsets and inflammation.

## Discussion

In recent years, there has been increasing recognition of the heterogeneity within the neutrophil population (43). However, our comprehension of neutrophil subsets remains relatively rudimentary, with subtle alterations observed in surface markers, gene expression profiles, and metabolic reprogramming. This study focused on identifying significant changes in surface protein expression among neutrophils across different tissue microenvironments, identifying a subset characterized by Stem cell antigen-1 (Sca1) expression. Stem cell antigen-1, Sca-1, belongs to the Ly-6 antigen family and is commonly used to identify murine hematopoietic stem cell (HSC) and progenitor populations (44) and also present on lymphocytes, myeloid cells to an extent (28, 45). Nonetheless, understanding Sca1 expression and its functional implications in neutrophils remains undefined. This previously undefined CD11b^+^Ly6G^+^Sca1^+^ or Sca1^pos^ neutrophil subset is distinct from earlier described monocytic cells exhibiting sca-1 expression in response to bacterial infection (28). This study sheds light on the potential roles of Sca1^pos^ neutrophils in tissue homeostasis and immune surveillance under steady-state and their association with inflammatory responses.

Given that Sca1 is not conventionally recognized as a neutrophil marker, pivotal efforts delve into the expression of Sca1 and its relevance to neutrophil morphology, phenotype, and functionality. Upon the revelation of Sca1 expression on CD11b^+^Ly6G^+^ neutrophils, we rigorously validated this discovery using a highly pure neutrophil population, defined as LiveCD45^+^CD11b^+^Gr1^hi^Ly6G^hi^Ly6C^lo^SiglecF^neg^CD115^-^ cells. Intriguingly, intracellular staining of Sca1 in neutrophils unveiled a substantial intracellular Sca-1 reservoir in BM and liver using flow cytometry and confocal microscopy. Transcript analysis provided additional support by validating the expression of the Ly6a gene in BM and liver neutrophils. These comprehensive findings firmly establish Sca1^pos^ neutrophils as a bona fide and distinctive subset within the neutrophil population.

Thorough investigations have pinpointed Sca1^pos^ neutrophils as a mature, activated, and inflammatory subset, apart from their Sca1^neg^ counterparts. This distinction is grounded on several compelling observations, including morphological features, phenotypic markers, and functional attributes. Sca1^pos^ neutrophils showcase hyper-segmented nuclei and heightened granularity, indicating their advanced maturation state. These neutrophils exhibit elevated expression levels of CD11b, LY6G, ICAM-1, CD47, CD49d, and CXCR4, typified markers for mature and activated cells (12, 46). Sca1^pos^ neutrophils demonstrate heightened ROS, NO, cytokines, enhanced phagocytic potential, and increased degranulation, indicative of their potent pro-inflammatory profile. Sca1^pos^ neutrophils also exhibit a notable accumulation of lipid droplets compared to their Sca1^neg^ counterparts. Lipid droplets in leukocytes contribute to inflammatory mediators such as eicosanoids (47). Moreover, they exhibit enhanced NET formation and T-cell proliferation. These characteristics align closely with those reported in other identified inflammatory neutrophil subsets, such as SiglecF^high^ and c-Kit^+^ neutrophils (20, 21). Furthermore, *in vitro* neutralization of Sca1 on neutrophils mitigated ROS generation and T cell proliferation, suggesting the functional relevance of Sca-1 on neutrophils, with ROS generated by neutrophils playing a pivotal role in modulating the inflammatory cell balance (48). Although Sca1 knockout mice do not exhibit significant alterations in hematological parameters and functions under steady-state conditions (49), further investigations for responses of Sca1 knockout mice concerning inflammatory contexts are warranted.

Nonetheless, our data refute that Sca1 expression on neutrophils is merely an ageing-related phenotype. This is based on no significant alterations in CD62L expression between Sca1^pos^ and Sca1^neg^ neutrophils, suggesting that CD62L regulation in neutrophils operates independently of Sca1 expression. However, some CD62L shedding was observed in liver tissue (50, 51). Adoptive transfer experiments indicated upregulation of Sca1 expression on the traced neutrophils in a tissue microenvironment-dependent manner rather than the chronological age of transferred cells. Lastly, similar levels of Sca1 expression on BrdU^neg/pos^ neutrophils suggest chronological ageing independent regulation of Sca1 expression on neutrophils.

Furthermore, comprehensive examinations elucidate the regulation of Sca1^pos^ neutrophils. Our findings reveal that Sca1 expression on neutrophils is upregulated in response to inflammatory stimuli, as evidenced upon exposure to various inflammatory cues *in vitro*. An increase in Sca1 expression in lineage-committed progenitors has been observed during septicemia, suggesting an inflammation-mediated induction of Sca1 expression (52). Moreover, the Toll-like receptor 4 (TLR4)/Sca-1 signaling axis via JNK signaling has been elucidated in bacterial infection-induced upregulation of Sca-1 during emergency myelopoiesis, particularly in myeloid progenitors (45). These findings underscore the significance of inflammatory stimuli in shaping the emergence of Sca1^pos^ neutrophils.

Conversely, tissue niche-dependent upregulation of Sca1 expression on neutrophils was also observed after adoptive transfer. BrdU labelling and intravascular staining of neutrophils further suggested the regulatory influence of prospective tissue environments on Sca1 expression. Further investigations are required to comprehend the appearance of Sca-1 on neutrophils, particularly in liver tissue. Notably, the adult liver typically does not express Sca-1 under normal circumstances, but its presence is observed in the fetal liver, presumably on hematopoietic cells (53). Moreover, during liver inflammation, murine hepatic endothelial and oval cells have been shown to express Sca1 (53, 54). These data suggest a nuanced role of inflammatory reprogramming and tissue milieu in regulating Sca1 expression on neutrophils.

There is a notable augmentation of Sca1^pos^ neutrophils in inflammatory settings, including peritonitis and NASH. The elevated frequency of inflammatory Sca1^pos^ neutrophils in NASH may contribute towards tissue damage, as suggested by the protective effect of neutropenia against MCD diet-induced steatohepatitis (55). Upregulation of Sca1 on myeloid macrophages has been associated with increased mortality in pathological conditions (28). Furthermore, antibody-mediated depletion of Sca1^pos^ cells has rescued survival defects and reduced tissue damage (28); the outcome might also depend on the likely depletion of Sca1^pos^ neutrophils. Still, it is technically challenging to deplete specific Sca1^pos^ neutrophils using antibody-mediated depletion concerning other Sca1-expressing cells. In this study, we observed the rescue of Sca1^pos^ neutrophils with NAC-dependent mitigation of acute inflammation (**Fig 6J**). However, the extent of their involvement in chronic inflammation necessitates further investigation. In addition to the liver and peritoneal cavity, Sca1^pos^ neutrophils were also enhanced in blood and BM in inflammatory settings. It will be interesting to understand the possibility of getting released into the circulation after their reprogramming concerning maturation in the liver, if any, under particular inflammatory stimulation.

Ly-6 gene superfamily has multiple members with different roles in neutrophil characteristics (56). Sca-1 expression is governed by the Ly6A/E molecule, which possesses two allelic variants, Ly6E and Ly6A. Certain mouse strains like C57BL/6, DBA/J and AKR/J express Ly6A while BALB/c, CBA/J and A/J strains express Ly6E (57). Our study confirmed LY6A/E expression in neutrophils from various tissues of C57BL/6 and BALB/c mice (**Fig S2A-C**). Ly6E expression has been documented on circulating monocytes during HIV infection (58). Moreover, it participates in restricting human coronavirus entry (59). A recent study using single-cell RNA-sequencing in pre-clinical tumor model predicted Ly6E^+^ neutrophils for immune checkpoint inhibitors (ICI) immunotherapy response (60). Additionally, recent studies have implicated the human ortholog Ly6A gene in pituitary tumors (61), while Ly6S is highly expressed by spleen cells and associated with inflammation (62). Furthermore, another member of the human Ly-6 gene superfamily, CD177 has garnered considerable attention in human neutrophils, with CD177^+^ neutrophils increasing in infection and inflammatory diseases (63, 64). These observations warrant further investigation into the immunological role of Ly6a and Ly6e expression on neutrophils across a spectrum of human health and disease.

## Conclusions

Our study identified and characterized Sca1^pos^ neutrophils, primarily abundant in liver tissue under steady-state conditions. We delineated the unique phenotypic and functional attributes of Sca1^pos^ neutrophils, highlighting their mature, activated, and pro-inflammatory nature. Furthermore, these neutrophils appear to develop through the *de novo* expression of Sca1 on Sca-1^neg^ neutrophils and are significantly increased in inflammatory contexts. Tracing experiments and the intravascular location of this cell have indicated that Sca1 expression on neutrophils is partly influenced by the liver tissue microenvironment, irrespective of age or inflammatory stimuli in homeostatic conditions. Furthermore, our study contributes to the broader understanding of neutrophil biology, providing insights into the potential roles of Sca1^pos^ neutrophils in inflammatory conditions, including NASH and peritonitis. Future investigations aimed at therapeutically targeting this neutrophil subset may offer insights into neutrophil-mediated inflammation.

## Experimental procedure

### Animals

The C57BL/6 and BALB/c male mice (eight to twelve weeks old) were used for the experiments. Mice were bred and kept in the animal house at CSIR-CDRI and fed with a chow diet. Until otherwise mentioned, data are provided from C57BL/6 mice. Mice were sacrificed at 9-10 AM to avoid any circadian rhythm-related variance in the study. All the experiments were performed per the IAEC approval guideline at CSIR-CDRI.

### Neutrophil isolation from mouse BM, blood, and different tissues

Different tissues like BM, blood, liver, lung, and spleen were collected after cardiac perfusion. BM, blood and spleen tissue cells were isolated without collagenase digestion. Neutrophils were isolated from BM and blood using Percoll gradient and Histopaque-1119/1077 gradient, respectively. Spleen cells were mechanically dissociated using a syringe plunger and further enriched with RBC lysis (10, 65). Liver and lung samples were finely minced and digested with 0.8mg/ml collagenase solution for 20 min at 37℃. Collagenase activity was stopped by transferring samples to ice and centrifugation at 4°C. To eliminate tissue clumps and debris, liver, and lung suspensions were centrifuged at differential speeds of 30 g for 3 min and 223 g for 10 min. The pelleted cells were re-suspended in HBSS 0.1%BSA and centrifugation at 850g for 30 min on a density gradient of Histopaque 1119 and 40% Percoll. Neutrophil-rich leukocytes were collected and washed (10). In some studies, BM and blood cells were isolated with collagenase treatment, similar to liver neutrophil isolation. The purity of neutrophils was evaluated on FACS ARIA Fusion after staining with CD45, CD11b, Ly6G, and DAPI for 20 min. Viability of neutrophils was always >95%, as assessed by trypan blue staining.

In some experiments, neutrophil enrichment was further achieved from blood and Liver with negative selection using B220, CD3e, Ter-119, F4/80, and CD5 antibodies (66). For the highest purity (>98%), CD11b^+^Ly6G^+^Sca1^+^ and CD1b^+^Ly6G^+^Sca1^-^ cells were sorted using FACS Aria Fusion (BD Biosciences) in PEB (PBS, 2mM EDTA, 10%FBS) buffer.

### Multi-color flow cytometry

The enriched leukocyte fractions of BM, blood, spleen, liver, and lung were stained with antibodies against the myeloid and neutrophil surface markers, including CD11b and Ly6G. DAPI or 7AAD was used for live/dead staining, single cell population was identified using SSC-A, SSC-H, and FSC-A, FSC-H gating strategy. Live single cells were analyzed in different combinations for various surface proteins using diverse antibodies in different colors gated for the CD11b^+^Ly6G^+^ neutrophil population among total leukocytes (**Fig S1A, Fig 1A, D, E**). These antibodies include FITC-CD11b, PE-Ly6G, Percp-CD45, PECy7-Sca1, APCCy7-c-Kit, BV605-CD115, BV421-CD19, PECF594-F4/80, APC-CD182(CXCR2), PE/Dazzel594-CD184(CXCR4), BV510-CD62L, APC-CD44, PE/Dazzle 594-CD54, BV421-CD47, Alexa fluor-CD49d, APCCy7-CD25, BV421-CD34, APC-B220, Percpcy5.5-CD11a/CD18, BV510-CD16/32, BV510-CD11c, APC-Cy7-Ly6C, Pacific blue-CD21/35. Samples were acquired on BD FACS Aria Fusion and analyzed using BD FACS Diva software.

For Intracellular staining of Sca1 in neutrophils, Single-cell suspensions isolated from the tissue were surface stained with CD11b, Ly6G, and 7AAD for 20 min. Following staining, cells were washed, fixed, and permeabilized with fixation and permeabilization buffer for 15 min in the dark. Cells were washed twice with 1x saponin wash buffer. Cells were stained with Sca1 for 20 min. Cells were washed, and samples were analyzed using BD FACS Diva software. After cell surface labelling, cells were washed and incubated with BODIPY 650/665 (2µM) for 15 min at 37°C in the dark to measure cellular lipid contents. Cells were washed and analyzed using a FACSARIA Fusion flow cytometer.

### Intravascular staining of neutrophils

In mice, 1 µg of Ly6G-PE antibody was injected intravenously. Blood was collected within 5 min, and the liver was perfused through the inferior vena cava. Liver-enriched leukocytes were surface stained with Percpcy5.5 CD45, FITC CD11b, APCCy7 Ly6G, PECy7 Sca1, live/dead dye DAPI and analyzed using a FACSARIA Fusion flow cytometer (67).

### Real-Time PCR

RNA was isolated from sorted Sca1^neg/pos^ neutrophils from BM and liver using the RNAesy mini kit (Quiagen) per manufacturer instructions. cDNA was prepared using a High-capacity cDNA Reverse Transcription Kit (Applied bioscience). Primers used for Ly6a (Forward CCTACCCTGATGGAGTCTGTGT, Reverse CACGTTGACCTTAGTACCCAGG) and Ly6e gene (Forward CCTGATGTGCTTCTCATGTACCG, Reverse GTTCAGGGTGTAGCCAAGGTTG). The relative fold differences between the groups were calculated by using the comparative cycle threshold (2^-ΔΔCt^) method, and actin mRNA was used as an internal control to calculate the relative mRNA expression.

### Giemsa staining

Isolated leukocytes or sorted neutrophil subsets (2 ×10^4^ cells) from different tissues were centrifuged at 500g for 5 min on slides. Cells were fixed with a May-Grunwald stain for 20 min, washed, and stained with Giemsa stain (1:10 dilution) for 30 min. Slides were washed, dried, and mounted with DPX mounting medium. Images were captured and analyzed using a Nikon Eclipse TS2 microscope with a 60× objective.

### Confocal microscopy

Sca1^neg^ and Sca1^pos^ neutrophils were sorted from BM, blood, and liver. Cells were adhered on a fibrinogen coated microwell chamber slide and fixed with 2% paraformaldehyde for 20 min. Cells were permeabilized with 0.1% triton-x for 20 min, washed, and blocked with 2%BSA for 20 min. Cells were stained with primary antibodies for Neutrophil Elastase (1:200) or Sca1 (1:50) overnight at 4 °C. The next day, cells were washed thrice and stained with secondary antibody Alexa Fluor 488 (1:500) and Rhodamine Phalloidin (1:200) for 2 h in the dark. Cells were washed thrice, mounted with a DAPI-containing mounting medium, and covered with a cover slip. Cells were observed using Leica DMI6000 microscope 40x and an Olympus confocal microscope with 60x (31, 65).

For lipid droplet staining, sorted cells were treated with 2µM Bodipy at 37 °C in the dark for 15 min. Cells were washed and fixed with 2% paraformaldehyde for 15 min. After washing, cells were stained with DAPI containing mounting medium. Cells were observed using a Leica DMI6000 microscope with 40x and an Olympus confocal microscope with 60x.

Liver tissue cryosections were fixed with chilled Acetone, permeabilized with 0.3% Triton X, and blocked with 3%BSA. Sections were stained overnight with primary antibody for NE (1:100) and Sca1 (1:100) at 4 °C. The next day, sections were washed, and secondary antibodies Alexa fluor 568 (1:400) and Alexa fluor 488 (1:400) for 2 h in the dark. Sections were washed and stained with a mounting medium containing DAPI. Images were captured using Leica DMI6000 microscope 40x.

### Adoptive transfer and BrdU labeling of neutrophils

Purified BM neutrophils (>99%Sca1^neg^) were labeled with 2μM CFSE dye for 15 min. 5 ×10^6^ cells were injected in mice intravenously. 16 h later, different tissues like BM, blood, lung, and liver were collected. Tissue single-cell suspension or enriched leukocyte fraction was surface stained for APCCy7-Ly6G, PECy7-Sca1, and DAPI. Samples were acquired by FACSARIA Fusion and analyzed for Sca1 expression on CFSE-labeled neutrophils.

BrdU (2.5 mg/kg) was injected intraperitoneally in mice. Tissues, including BM, blood, lung, and liver, were collected at 72 h. Isolated cells from different tissues were surface stained with PE-CD11b, APCCy7-Ly6G, PECy7-Sca1, and DAPI. Cells were fixed and permeabilized with a fix and permeabilization buffer. Cells were treated with 30 µg/ml DNase for 1h at 37 °C and stained with anti-BrdU antibody for 20 min. Samples were acquired on BD FACSARIA Fusion (15, 18).

### Measurement of Superoxide, Nitric oxide, Mitochondria, and mitochondrial superoxide

BM, blood, and liver leukocytes were stained with CD11b, Ly6G, and Sca1 antibodies and live/dead dye, 7AAD, at 4 °C. For superoxide and nitric oxide detection, cells were stained using DHE (10µM), and DAF-2DA (2µM) for 30 min. Mitochondrial membrane potential and superoxide were detected with Mito tracker Red (50nM) and MitoSox (2µM) for 30 min. Cells were washed, and samples were acquired on FACSARIA Fusion (31, 65).

### *In vitro* blockade of Sca1

To analyze the functional relevance of Sca1 on neutrophils, isolated liver neutrophils were blocked with 10 µg/ml α-Sca1 antibody clone E13-161.7 *in vitro* (28). Superoxide levels were detected with/without PMA stimulation by FACS (28).

### Phagocytosis Assay

Isolated neutrophils from BM and enriched neutrophils with a negative selection from blood and liver were surface stained with CD11b, Ly6G, Sca-1 and DAPI for 20 min. PE-labeled latex beads were opsonized with 10% FBS for 30 min and centrifuged at 10,000g. Neutrophils were co-cultured with beads at Neutrophil: bead ratio of 1:10 in 48 well flat bottom plates in RPMI medium containing 5% FBS at 37 °C for 90 min. In other studies, *E. coli* were tagged with 2mg/ml FITC, washed thrice with HBSS-5% FBS for 10 min at 10000g. Bacteria were cocultured with neutrophils with 1 MOI and 5 MOI for 15 min. After incubation, cells were centrifuged at 200g for 5 min. Samples were acquired using BD FACSARIA Fusion flow cytometer (68).

### Electron microscopy and Degranulation assay

Sorted Sca1^neg^ and Sca1^pos^ neutrophils were fixed with 2.5% glutaraldehyde at RT for 1h and transferred to 4°C overnight. The next day, cells were washed and stained with Evans Blue (2mg/ml). After washing, the blue color cell pellet was resuspended in 2% low melting agarose. Cells were centrifuged and kept on ice for 15 min to solidify the agar. The pellet was scraped and treated with 1% OsO4 solution for 2 h at RT and processed with an ascending range of alcohol. Following this, cells were treated with an ascending range of acetone and embedded in plastic resin overnight. Ultrathin sections (∼70 nm) were cut using a Leica EM UC7 ultramicrotome, collected on copper grids, and stained with lead citrate and uranyl acetate. Images were captured with JEOL JEM1400 electron microscope (69).

For degranulation analysis, control and PMA-stimulated BM, blood, and liver leukocytes were stained with CD11b, Ly6G, Sca1, 7AAD, and CD63 (primary granule) 30 min. Cells were washed, and %Sca1^neg/pos^ CD63^+^ neutrophils were analyzed on BD FACSARIA Fusion.

### NETs releasing ability and Histone citrullination

To check ROS dependent or independent NETosis, sorted Sca-1^neg/pos^ from neutrophils BM and liver were on fibrinogen coated microwell chamber slide with/without PMA and ionomycin treatment for 3h. Cells were fixed, permeabilized, and stained for neutrophil elastase primary antibody (1:200). Cells were washed and stained with anti-rabbit Alexa Fluor 488 secondary antibody for 2h at RT. After washing, slides were mounted with DAPI containing mounting medium. NETs were identified as decondensed nuclei, mixing with granular protein elastase and release of the same. Images were captured on a Leica DMI6000 microscope with 40x.

Isolated leukocytes from BM, Blood, and liver were stained for CD11b, Ly6G, and Sca1 surface staining with live/dead staining of 7AAD. An H3Cit antibody followed by a secondary antibody was used to detect H3Cit expression. Cells were washed and analyzed by FACSARIA Fusion.

### T-cell proliferation assay

Splenic T cells were stained with CFSE (1μM, 5min at RT in dark), activated with soluble anti-CD3e and anti-CD28 antibody, and co-cultured with Sca-1^neg/pos^ neutrophils sorted from BM and liver in RPMI medium containing 5%FBS in 96 well plates. T cell: PMN ratio was 1:1, 1:3 or 1:5. After 48h, CFSE dilution was observed by flow cytometry to check T cell proliferation. In some experiments, Sca1^neg/pos^ neutrophils and T cells were co-cultured in the presence of N-acetyl cysteine (10 mM), Mito Tempo (10 µM) and α-Sca-1 antibody. After 48h, CFSE dilution was observed by flow cytometry.

### Intracellular cytokine measurement

Leukocytes isolated from different tissues were treated with LPS (1µg/ml), followed by Brefeldin A (10μg/ml) for 3 h at 37 C° to retain cytokine inside cells. After incubation, cells were surface stained with CD11b, Ly6G, Sca1, and 7AAD. Cells were washed and treated with fixation and permeabilization buffer for 15. Cells were washed twice with 1x saponin wash buffer and stained with fluorescent conjugated anti-cytokine antibodies like APC-anti-IL6 and AlexaFluor488 IL1β (28). Intracellular cytokine levels were analyzed with respect to FMO control on BD FACSARIA Fusion.

### *In vitro* stimulation of conventional neutrophils and GMPs for Sca-1^neg/pos^ neutrophil switching

Purified BM neutrophils (>99% Sca-1^neg^) were cultured for 16 h in RPMI medium in presence of various inflammatory mediators like fMLP (5µM), PMA (10nM), Zymosan (10µg/ml), LPS (1µg/ml), ATP (100µM), LPS+ATP, at 37 °C. Cells were stained with CD11b, Ly6G, Sca1, and 7AAD and Sca-1^neg/pos^ neutrophils were analyzed by flow cytometry. An *in-vitro* granulopoiesis assay using progenitors GMPs in the absence/presence of inflammatory stimuli was used, and the extent of Sca-1^pos^ neutrophil production was determined (28). GMPs were differentiated in IMDM+10% FBS + SCF and TPO (200 ng/ml) in the presence or absence of G-CSF (100 ng/ml). On day 2, modulators LPS (20ng/ml) and Palmitic Acid (5 µg/ml) were added. On day 6, cells were stained with CD11b, Ly6G, Sca1, and DAPI to analyze the Sca-1 expression by flow cytometry.

### Inflammation model in animals

For *in vivo* peritonitis inflammation, mice were given 2 mg/kg LPS or PBS via i.p. After 16 h, mice were sacrificed, and BM, blood, and peritoneal fluid were analyzed for Sca1 expression on neutrophils. In some studies, mice were pretreated with 200 mg/kg/day NAC for three constitutive days. On 4^th^ day, after 1 h of NAC pretreatment, LPS or PBS was injected intraperitoneally., mice were sacrificed after 16 h, leukocytes and neutrophil subsets were analyzed in BM, blood, peritoneal fluid, and liver by BD FACSARIA Fusion.

For the Diet-induced NASH model, C57BL/6 mice were fed a chow diet or methionine and choline-deficient (MCD) diet. Systemic serum AST ALT levels and liver/body weight ratio were monitored as indicators of NASH. An EM 200 auto-analyzer (Erba, Transasia) was used to compute the serum ALT and AST activity by the test kit instructions. Liver tissues from CD and MCD-fed mice were fixed in a 10% formalin. 5 μm thick paraffin sections were cut from formalin-fixed paraffin-embedded tissue blocks, deparaffinized, rehydrated, and stained with eosin and hematoxylin or picrosirius red. The slides were visualized under a light microscope.

### Statistical analysis

All experiments were performed 3-8 times. Mice were randomly assigned to different experimental groups. The group sizes (n), including biological or technical replicates, specific statistical tests used to determine significance, and p values are provided in the figure legends. P value < 0.05, significant. All statistical analyses were performed using Prism (GraphPad).

## Author contributions

M.N. performed, analyzed experiments, interpreted data, prepared figures, and wrote the manuscript. S.S., P.D., A.S., and K.G. performed and analyzed specific experiments. A.S. and A.G. helped with the NASH animal model, and K.J. supervised the NASH study and provided critical input on the manuscript. R.S. and K.M. performed and provided critical input on TEM analyses. A.L. and M.D.F. analyzed and interpreted experiments and provided crucial support. M.D. contributed reagents and provided support for both experimental design and analysis. S.K. conceptualized, designed, and interpreted experiments, reviewed the progress, and finalized the manuscript.

## Acknowledgements

We thank Mr. AL Vishwakarma at Flow cytometry Core for his help with flow cytometry experiments. We also acknowledge DMI6000 facility at the pharmacology division and the Olympus confocal microscope at the Core facility. We thank the Director CSIR-CDRI for the support during the study. We acknowledge the financial support of the early career grant from DST-SERB ECR/2017/001274 and CRG/2022/001939. MD is supported by SERB (JBR/2020/000034). All significant data generated or analyzed during this study are included in this article, and any specific data are available upon reasonable request from the corresponding author. This manuscript is the CDRI communication number <u>50/2024/SK.</u>

