## Supplementary Information for "Sca-1 expression depicts pro-inflammatory murine neutrophils under steady state and pathological conditions"

**Supplementary Fig 1**

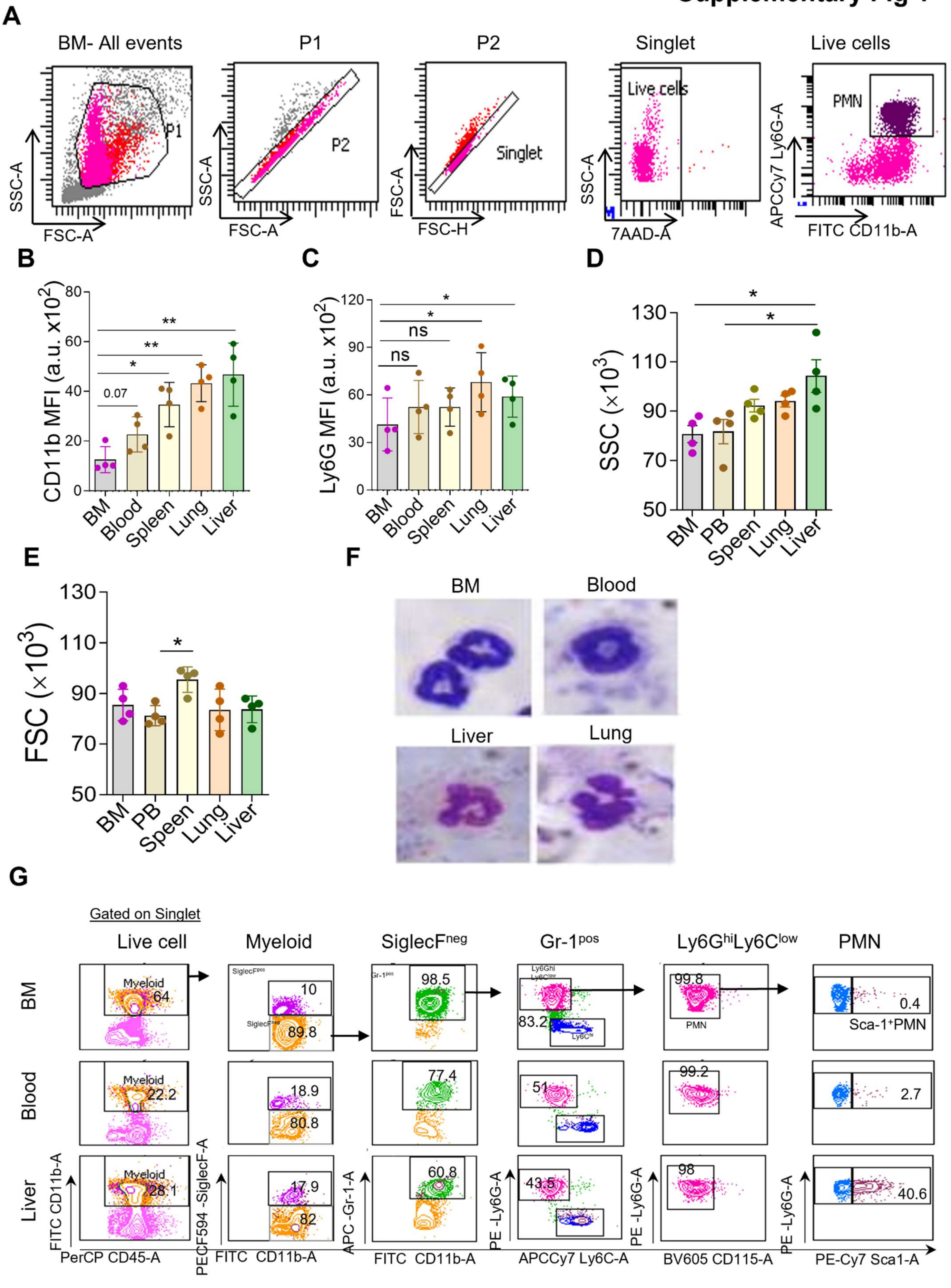

**Supplementary figure1:**

**Gating strategies for neutrophils from different tissues and their characteristics**

**A.** Gating strategies for Flow cytometric analysis of $7AAD^{-}CD11b^{+}Ly6G^{+}$ live neutrophils. n=4 biological replicates.

**B.** Bar graph represents the quantification of CD11b MFI from Flow cytometric data. N=4 biological replicates. *$p<0.05$, **$p<0.01$ using Student's paired t-test.

**C.** Bar graph represents the quantification of Ly6G MFI from Flow cytometric data. n=4 biological replicates. *$p<0.05$ using Student's paired t-test.

**D.** Bar graph represents the quantification of SSC mean from Flow cytometric data. n=4 biological replicates. *$p<0.05$, using Student's paired t-test.

**E.** FSC data presented as bar graph for neutrophils isolated from different tissues. n=4 biological replicates. *$p<0.05$, using Student's paired t-test.

**F.** Giemsa staining represents the nuclear morphology of different tissue-derived neutrophils. n=4 biological replicates.

**G.** Gating strategy represents elimination of minor contamination of $SiglecF^{+}$ eosinophil and $Ly6C^{hi}CD115^{+}$ monocyte cells in Singlet $7AAD^{-}$ live $CD45^{+}CD11b^{+}$ myeloid cells presenting Sca-1 expression on highly pure $CD45^{+}CD11b^{+}Ly6G^{hi}Ly6C^{low}$ neutrophils in BM, blood and liver. Data represents n=4 biological replicates.

### Supplementary Fig 2

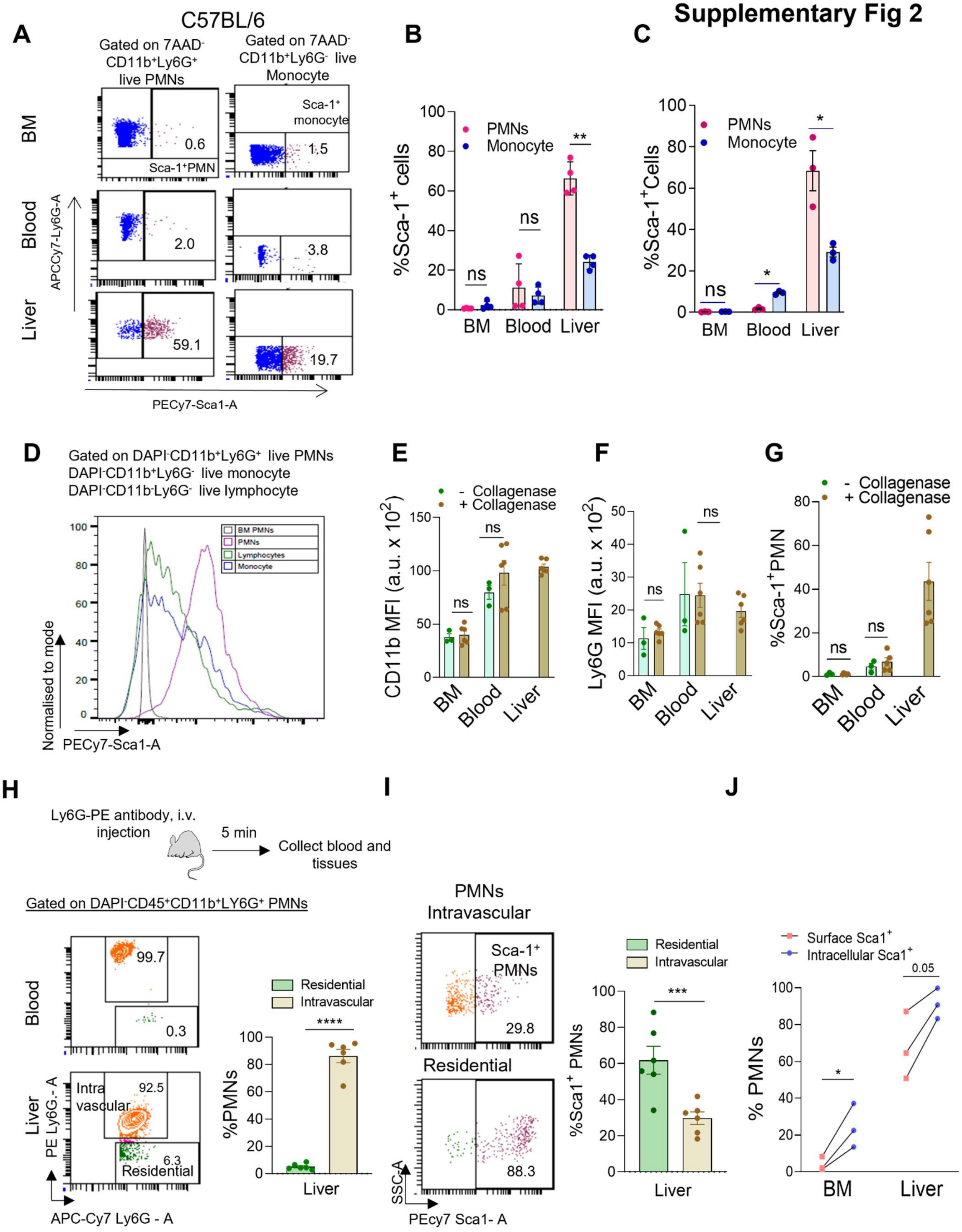

**Supplementary figure 2:**

**Sca1⁺ PMNs on different tissues of C57BL/6 and BALB/c mice**

**A.** Flow Cytometric dot plot represents %Sca1$^+$ neutrophils (CD11b+Ly6G+Sca1$^+$) and %Sca1$^+$ monocyte (CD11b+Ly6G$^-$Sca1$^+$) of BM, blood and liver of C57BL/6 mice. Data represents n=4 biological replicates.

**B.** Quantification of %Sca1$^+$ cells (PMNs and monocytes) in BM, blood, and liver of C57BL/6 mice. n=3-4 biological replicates. **p<0.01 using Student's paired t-test.

**C.** Quantification of %Sca1$^+$ cells (PMNs and monocytes) in BM, blood, and liver of BALB/c mice. n=3-4 biological replicates. *p<0.05 using Student's paired t-test.

**D.** Histogram overlay shows Sca1 expression on PMNs, lymphocytes, and monocytes in liver tissue of C57BL/6 mice with reference to BM PMNs. Data is representative of n=4 biological replicates.

**E.** Bar graph represents the quantification of CD11b MFI from flow cytometric data of BM, blood, and liver PMNs with/without *in vitro* collagenase treatment. N= 3-6 biological replicates. ns= non-significant by Student's unpaired t-test

**F.** Bar graph represents the quantification of Ly6G MFI from flow cytometric data of BM, blood, and liver PMNs with/without *in vitro* collagenase treatment. N= 3-6 biological replicates. ns= non-significant by Student's unpaired t-test

**G.** Bar graph represents the quantification of %Sca1$^+$ PMNs with/without *in vitro* collagenase treatment to BM, blood, and liver PMNs. Data represents n= 3-5 biological replicates. ns= non-significant by Student's unpaired t-test

**H.** Experiment strategy for intravascular labeling of neutrophils**.** Flow cytometry density plot represents intravascularly labeled vs resident neutrophils of blood and liver and its quantification of liver tissue. Data represents n= 6 biological replicates ****p<0.0001 using Student's unpaired t-test.

**I.** Flow cytometry dot plot represents %Sca1$^+$ PMNs of liver intravascular vs resident neutrophils and its quantification. Data represents n= 6 biological replicates ***p<0.001 using Student's unpaired t-test.

**J.** Flow cytometry data quantification of the percentage of intracellular vs. surface Sca1 expression on BM and liver neutrophils. Data are representative of n=3 biological replicates. *p<0.05 by Student's paired t-test.

All data are mean ± SE.

### Supplementary Fig 3

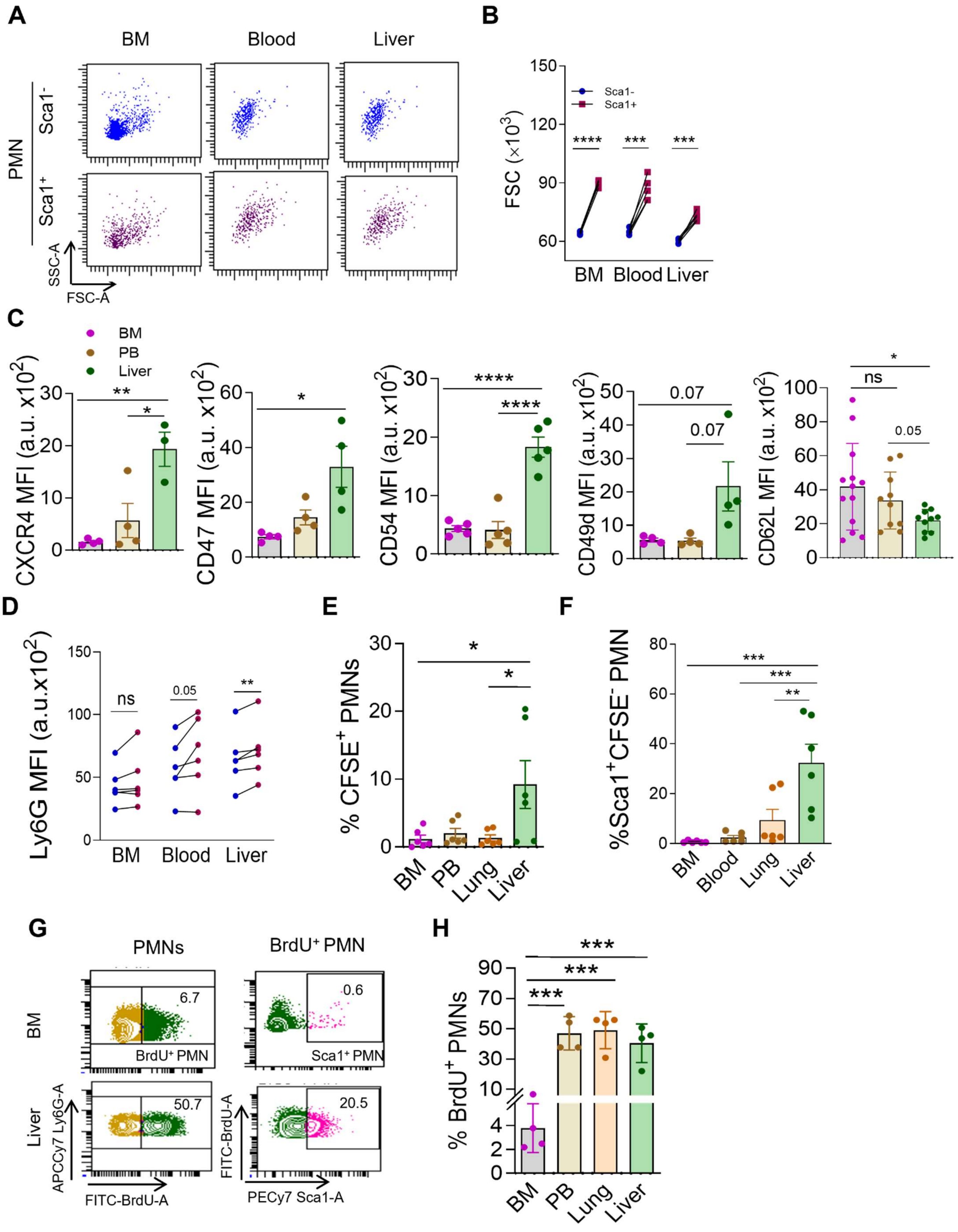

**Supplementary figure 3:**

**FSC and SSC analysis of $Sca1^{-/+}$ neutrophils, cell surface marker analysis of tissue neutrophils and CFSE and BrdU labeling of neutrophils in different tissues**

**A.** Dot plot representative of FSC and SSC of $Sca1^{-/+}$ neutrophils of BM, blood and liver. n=5 biological replicates.

**B.** Flow cytometry-based quantification of FSC of $Sca1^{-/+}$ neutrophils of BM, blood and liver. n=5 biological replicates. ***p<0.001, ****p<0.0001 by Student's paired t-test.

**C.** Bar graphs represent quantification of CXCR4, CD47, CD54, CD49d, CD62L on BM, blood and liver neutrophils. n=3-4 biological replicates, *p<0.05, **p<0.01, ****p<0.0001 by Student's paired t-test.

**D.** Quantification of Ly6G MFI on $Sca1^{-}$ and $Sca1^{+}$ neutrophils of BM, blood and liver. n=5 biological replicates. **p<0.01, ***p<0.001 using Student's paired t-test).

**E.** Flow cytometry-based quantification of $\%CFSE^{+}$PMNs of different tissue. n=6 biological replicates. *p<0.05 using One way ANOVA.

**F.** Bar graph represents $\%Sca1^{+}CFSE^{-}$ PMNs. n=6 biological replicates. **p<0.01, ***p<0.001 using One way ANOVA.

**G.** Contour plots represent $\%BrdU^{+}$ PMNs as well $\%BrdU^{+}Sca1^{+}$ PMNs in BM and liver. n=4 biological replicates

**H.** Quantification of $BrdU^{+}$ PMNs in different tissues analyzed ***<0.0001, by One way ANOVA test.

All data are mean ± SE.

**Supplementary Fig 4**

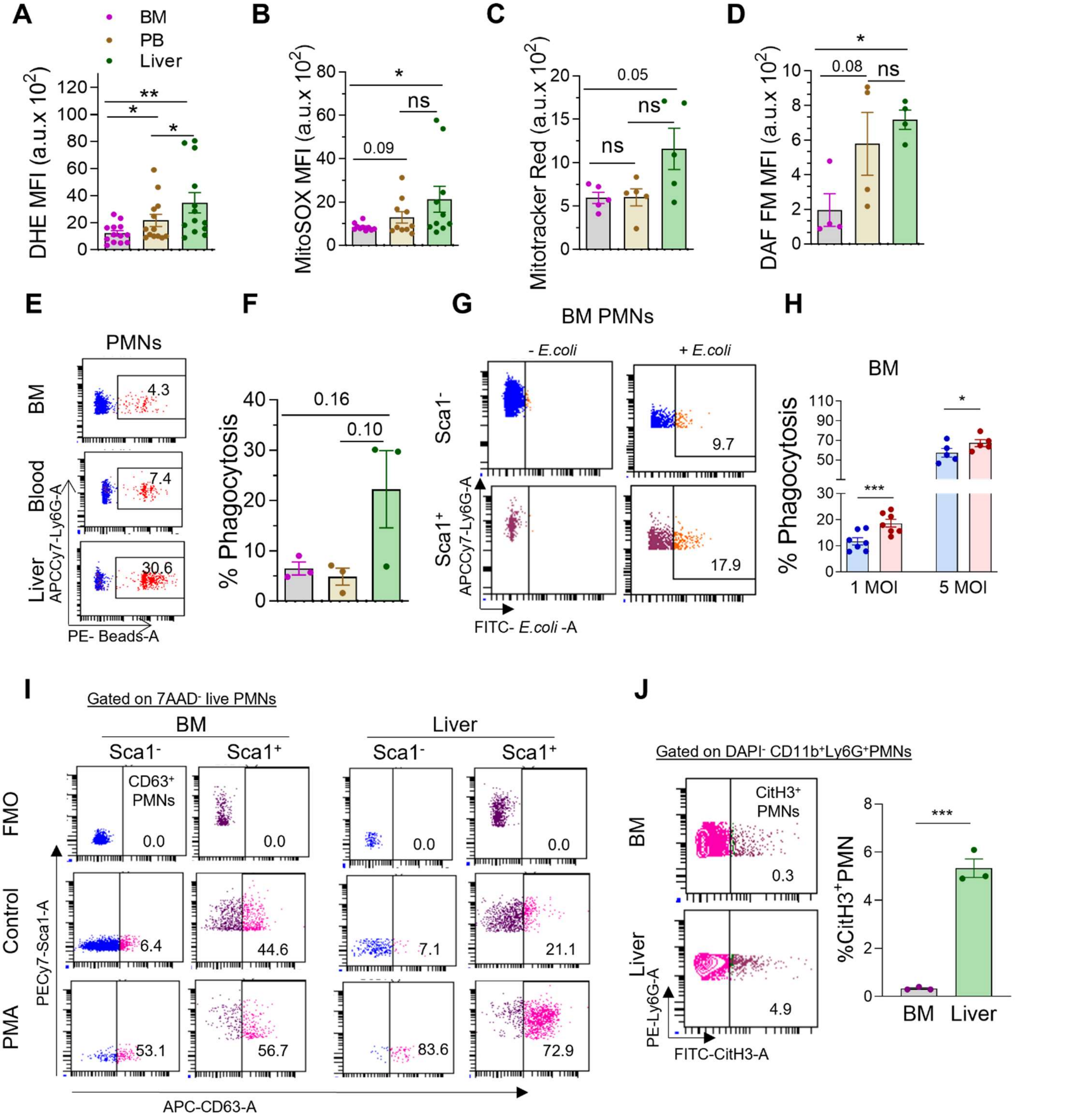


**Supplementary figure 4:**

**Analysis of functions of different tissue neutrophils and Sca1$^{-/+}$ neutrophils,**

**A.** Flow cytometry-based quantification of superoxide anion detected by DHE in tissue neutrophils. n=10-12 biological replicates; *p<0.05, **p<0.01 using Student's paired t-test.

**B.** Mitochondrial superoxide by MitoSOX staining in tissue neutrophils. n=10 biological replicates. *p<0.05 using Student's paired t-test.

**C.** Analysis of Mito tracker Red in tissue neutrophils. n=5 biological replicates; Exact p-value mentioned between marked groups, using Student's paired t-test.

**D.** Nitric oxide (NO) level detected by DAF FM in tissue neutrophils. n=3 biological replicates. *p<0.05, using Student's paired t-test.

**E.** Flow cytometry dot plot represents %phagocytosis of latex beads by different tissue neutrophils. Data represent n=3 biological replicates.

**F.** Quantification of %Phagocytosis of latex beads by different tissue neutrophils. n=3 biological replicates. Exact p-value mentioned between marked groups, using Student's paired t-test.

**G.** Flow cytometry dot plot represents %phagocytosis of *E. coli* by BM neutrophils at 1MOI. Data represent n=5-7 biological replicates.

**H.** Quantification of %phagocytosis of *E. coli* by BM neutrophils at 1MOI and 5MOI. Data represent n=5-7 biological replicates. *p<0.05, ***p<0.001 using Student's paired t-test.

**I.** Dot plots represent CD63 expression on $Sca1^-$ and $Sca1^+$ PMNs of BM and Liver with/without PMA treatment. n=4 biological replicates.

**J.** Contour plot represents $\%CitH3^+$ PMNs in BM and Liver and its quantification. n=3 biological replicates. ****p<0.0001 using Student's paired t-test.

All data are mean ± SE.

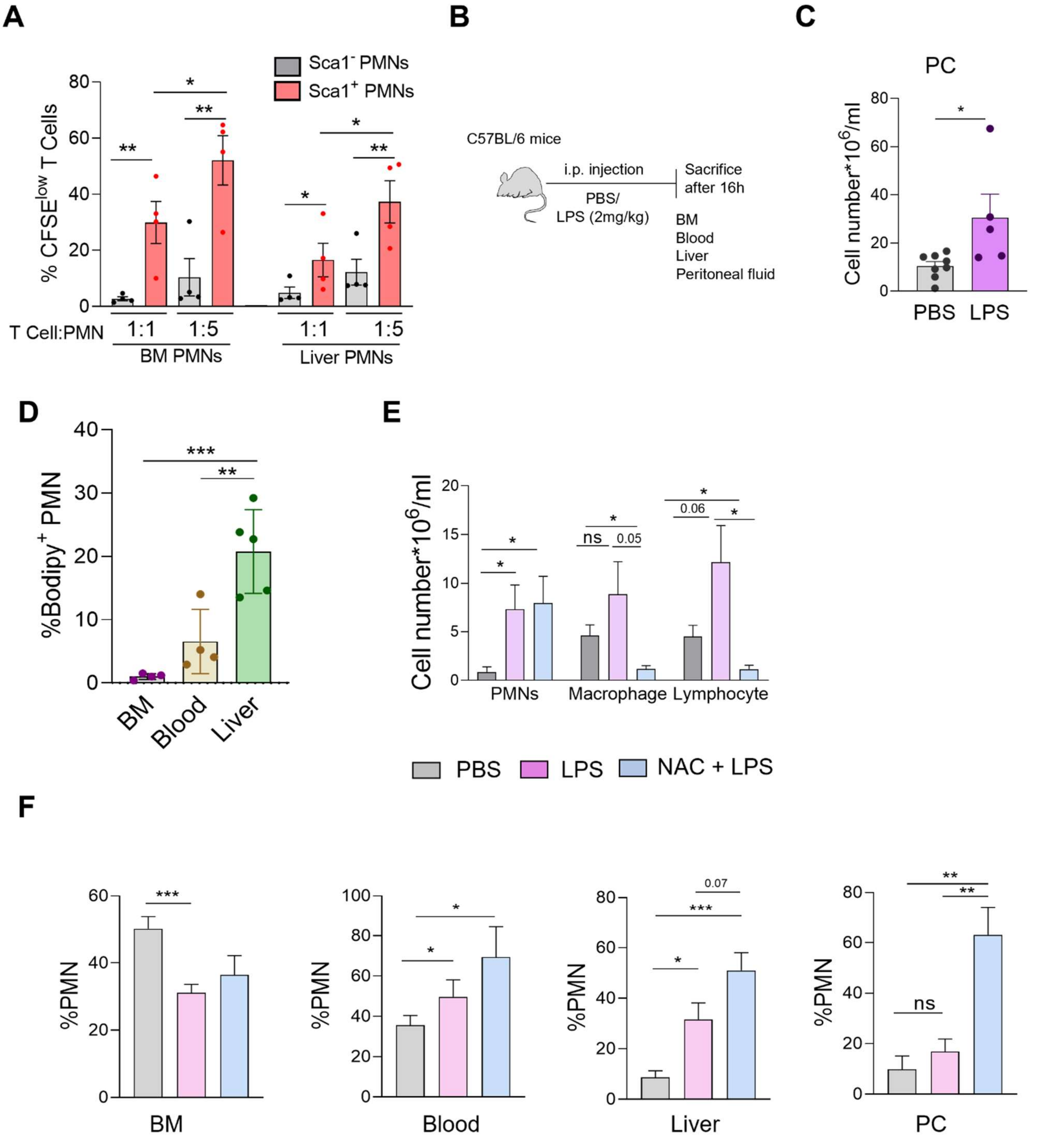


**Supplementary figure 5:**

**T cell proliferation by Sca1$^{-/+}$ PMNs and enrichment of Sca1$^{+}$ PMNs in density-based fractions**

**A.** Quantification of CFSE$^{low}$ T cell presenting division after coculture with Sca1$^{-}$ and Sca1$^{+}$ neutrophils with T cell: PMNs ratio of 1:1 and 1:5. n=4 biological replicates. *p<0.05, **p<0.01, using Student's paired t-test.

**C.** Experiment setup of LPS induced peritonitis model and quantification of total cells number of peritoneal cavity in control and LPS treated mice. n=5-7 biological replicates. *p<0.05 using Student's paired t-test.

**D.** Bar graph represents % Bodipy$^+$ PMNs in BM, Blood and Liver neutrophils. n=4-5 biological replicates. *p<0.05, **p<0.01, ***p<0.001 using Student's paired t-test.

**E.** Quantification of cell number of PMNs, macrophage and lymphocyte of peritoneal cavity in control and LPS treated mice with/without NAC treatment. Data represent n=4-6 mice.

**F.** Quantification of %PMNs in BM, Blood, Liver, and peritoneal cavity of control and LPS-treated mice with/without NAC treatment. Data represent n=2 experiments with 6-8 mice. *p<0.05, **p<0.01, ***p<0.001, using Student's paired t-test.

All data are mean ± SE.

| Reagent/Resource | Reference or Source | Identifier/ Catalog No. | Clone |
|---|---|---|---|
| **Experimental Models** | | | |
| C57BL/6J (*M. musculus*) | CSIR CDRI Animal facility | N/A | N/A |
| *BALB/c* | CSIR CDRI Animal facility | N/A | N/A |
| **Mice diet** | | | |
| Chow diet | Altromin, Germany | 1320 | N/A |
| MCD diet | Research Diets, Inc. | A02082002BR | N/A |
| **Antibodies** | | | |
| CD45- Alexa Fluor700 | Biolegend | 103128 | 30-F11 |
| CD45-PerCP- Cy5.5 | BD Pharmingen | 550994 | 30-F11 |
| CD11b- FITC | Biolegend | 101206 | M1/70 |
| CD11b- PE | Biolegend | 101208 | M1/70 |
| Ly6G/Ly6C(Gr-1) | Biolegend | 108448 | RB6-8C5CD |
| Ly6G- PE | BD Pharmingen | 551461 | 1A8 |
| Ly6G- APCCy7 | BD Bioscience | 560600 | 1A8 |
| Ly6G-Biotin | Biolegend | 750000264 | 1A8 |
| F4/80- PE-CF594 | BD Bioscience | 565613 | T45-2342 |
| SiglecF- PE-CF594 | BD Bioscience | 562757 | E50-2440 |
| CD19-BV421 | Biolegend | 115538 | 6D5 |
| CD19-V450 | BD Horizon | 560375 | 1D3 |
| Sca1(Ly6A/E)-PEcy7 | BD Pharmingen | 558162 | D7 |
| Purified Ly6A/E (Sca1) | Biolegend | 108102 | E13-161.7 |
| CD49d- Alexa Fluor 647 | Biolegend | 103614 | R1-2CD5 |
| CD16/32- PerCP-Cy5.5 | BD Bioscience | 560540 | 2.4G2 |
| CD16/32- BV510 | BD OptiBuild | 740111 | 2.4G2 |
| CD44-APC | BD Pharmingen | 559250 | IM7 |
| CD21/35-Pacific Blue | Biolegend | 123414 | 7E9 |
| CD34- FITC | BD Pharmingen | 553733 | RAM34 |
| CD34 PE | BD Pharmingen | 551387 | RAM34 |
| CD34-BV421 | BD Horizon | 562608 | RAM34 |
| CD115-BV605 | Biolegend | 135517 | AFS98CD11 |
| CD3e-PECy7 | Biolegend | 100220 | 17A2 |
| CD3e-Biotin | BD Pharmingen | 553060 | 145-2C11 |
| CD28-Biotin | BD Pharmingen | 553296 | 37.51 |
| CD11c-BV510 | BD Horizon | 562949 | HL3 |
| CD62L-BV510 | BD Bioscience | 563117 | MEL-14 |
| CD62L- PECy7 | BD Bioscience | 560516 | MEL-14 |
| CD117-APC | BD Pharmingen | 553356 | 2B8 |
| CD117-PerCP-Cy5.5 | BD Pharmingen | 560557 | 2B8 |
| CD182(CXCR2)-APC | Biolegend | 149312 | SA044G4 |
| CD184(CXCR4)-PEDazzel-594 | Biolegend | 146514 | L276F12 |
| CD47-BV421 | Biolegend | 127527 | miap301 |
| CD54- PE/Dazzel-594 | Biolegend | 116130 | YN1/1.7.4 |
| Ly6C-APCCy7 | BD Bioscience | 560596 | AL-21 |

| | | | |
|---|---|---|---|
| Biotin Mouse Panel | BD Pharmigen | 559971 | N/A |
| Biotin anti-F4/80 | Biolegend | 123106 | BM8 |
| Biotin anti-CD5 | Biolegend | 100604 | 53-7.3 |
| Annexin V- APC | Biolegend | 640920 | N/A |
| CD63-APC | Biolegend | 143905 | NVG-2 |
| IL-6- APC | Biolegend | 504507 | MP5-20F3 |
| IL1-β-Alexa Fluor 488 | Santacruz | SC12742 | N/A |
| Anti-neutrophil elastase antibody | Calbiochem | 481001 | N/A |
| Anti-Histone H4 (Citrulline H3) | Merck | 07-596 | N/A |
| Streptavidin-APC-Cy7 | BD Bioscience | 554063 | N/A |
| Streptavidin BV605 | BD Bioscience | 563260 | N/A |
| Donkey anti rat IgG Alexa Fluor 488 | Invitrogen | SA5-10026 | N/A |
| Donkey anti mouse IgG Alexa Fluor 488 | Invitrogen | A21202 | N/A |
| Donkey anti rabbit IgG Alexa Fluor 568 | Invitrogen | A10042 | N/A |
| Donkey anti rabbit IgG Alexa Fluor 488 | Invitrogen | A21206 | N/A |
| **Cytokines** | | | |
| Recombinant murine SCF | Biolegend | 579702 | N/A |
| Recombinant murine TPO | Biolegend | 593302 | N/A |
| G CSF | PROSPEC | cyt-410-b | N/A |
| **Dyes, inhibitor, other reagents** | | | |
| 7AAD | Invitrogen | A1310 | N/A |
| DAPI | Sigma | D9542 | N/A |
| DCF-DA | Sigma | D6883 | N/A |
| MitoSox-Red | Invitrogen | M36008 | N/A |
| DHE-PE | Cayman | 12013 | N/A |
| DAF-FM diacetate | Cayman | 18767 | N/A |
| Mitotracker red | Molecular probes | M7512 | N/A |
| BODIPY 650/665 | Invitrogen | D10001 | N/A |
| CFSE | Invitrogen | C34570 | N/A |
| N-Acetyl cysteine | TCI | A0905 | N/A |
| Mito Tempo | Cayman | 16621 | N/A |
| Sytox Green | Invitrogen | S7020 | N/A |
| Triton X-100 | Sigma | T9284 | N/A |
| $H_2O_2$ | Sigma | 323381 | N/A |
| Acetone | SRL | 15168 | N/A |
| Fluoromount- Mounting Medium, with DAPI | Thermo scientific | 00-4959-52 | N/A |
| DPX mounting media | Merck | DA5DF41649 | N/A |
| LPS | Sigma | L3012 | N/A |
| fMLP | Sigma | F3506 | N/A |
| Zymosan A | Cayman | 21175 | N/A |
| PMA | Cayman | 10008014 | N/A |
| Ionomycin | Cayman | 10004974 | N/A |
| Palmitic acid | Sigma | P0500 | N/A |
| Butyrate | Sigma | 303410 | N/A |
| Oleic acid | Cayman | 90260 | N/A |
| ATP | Cayman | 14498 | N/A |

| | | | |
|---|---|---|---|
| 5-bromo-2'-deoxyuridine (BrdU) | BD bioscience | 51-2420KC | N/A |
| Brefeldin A | Cayman | 11861 | N/A |
| Tissue freezing media | Sigma | SHH0026 | N/A |
| Poly-L-Lysine | Sigma | P8920 | N/A |
| Fibrinogen | Sigma | F8630 | N/A |
| Paraformaldehyde | Sigma | P6178 | N/A |
| Osmium tetra oxide | Sigma | 201030 | N/A |
| Glutaraldehyde | Sigma | G5882 | N/A |
| BSA | GLR innovation | GLRIN19021909 | N/A |
| Hematoxylin | Sigma | H3136 | N/A |
| Eosin | Sigma | E4382 | N/A |
| Sirius red | Sigma | 365548 | N/A |
| Picric acid solution | Sigma | P6744 | N/A |
| May-Grunwald stain | HIMEDIA | S039 | N/A |
| Giemsa stain | HIMEDIA | TCL083 | N/A |
| Evans blue | Sigma | E-2129 | N/A |
| FBS | Gibco | 10270106 | N/A |
| EDTA | Sigma | E-6511 | N/A |
| Latex beads-PE | Sigma | L2778 | N/A |
| RPMI-1640 media | Lonza | 12-702F | N/A |
| Penstrep glutamine | Gibco | 1881463 | N/A |
| Percoll | Cytvia | 17089101 | N/A |
| 10X HBSS | Gibco | 14185052 | N/A |
| **Oligonucleotides and other sequence-based reagents** | | | |
| Ly6a primer | Thermo scientific | Mm00726565_s1 | N/A |
| Actin primer | TaqMan | Mm02619580_g1 | N/A |
| PCR master mix | Promega | M750B | N/A |
| **Software** | | | |
| Prism8.0 | GraphPad | https://www.graphpad.com | N/A |
| BD FACS Diva | BD Bioscience | https://www.bdbiosciences.com | N/A |
| FlowJo V.10 | BD Bioscience | https://www.flowjo.com/ | N/A |
| FV10-ASW viewer | Olympus | www.olympus-lifescience.com | N/A |
| Leica LAS AF | Leica | https://www.leica-microsystems.com/ | N/A |
| **Kits, glassware, plastic ware** | | | |
| High-capacity cDNA reverse transcriptase | Applied Bioscience | 4368814 | N/A |
| RNeasy Mini Kit | Qiagen | 74104 | N/A |
| MicroAmp Fast 96 well reaction plate (0.1mL) | Applied biosystem | 4346907 | N/A |
| Optical Adhesive Covers | Applied biosystem | 4360954 | N/A |
| 96-well U bottom plates | Genetix | 34296 | N/A |
| 48 – well flat bottom plate | Genetix | 32048 | N/A |
| 24- well flat bottom plate | Genetix | 32024 | N/A |
| 12- well flat bottom plate | Genetix | 32012 | N/A |
| 6- well flat bottom plate | Genetix | 32006 | N/A |
